# A mouse model to study glutathione limitation *in vivo*

**DOI:** 10.1101/2024.01.08.574722

**Authors:** Rebecca C. Timson, Artem Khan, Beste Uygur, Marwa Saad, Hsi-Wen Yeh, Nicole DelGaudio, Ross Weber, Hanan Alwaseem, Jing Gao, Chingwen Yang, Kıvanç Birsoy

## Abstract

Glutathione (GSH) is a highly abundant tripeptide thiol that performs diverse protective and biosynthetic functions in cells. While changes in GSH availability are linked to many diseases, including cancer and neurodegenerative disorders, determining the function of GSH in physiology and disease has been challenging due to its tight regulation. To address this, we generated cell and mouse models that express a bifunctional glutathione-synthesizing enzyme from *Streptococcus Thermophilus* (GshF). GshF expression allows efficient production of GSH in the cytosol and mitochondria and prevents cell death in response to GSH depletion, but not ferroptosis, indicating that GSH is not a limiting factor under lipid peroxidation. CRISPR screens using engineered enzymes revealed metabolic liabilities under compartmentalized GSH depletion. Finally, GshF expression in mice is embryonically lethal but sustains postnatal viability when restricted to adulthood. Overall, our work identifies a conditional mouse model to investigate the role of GSH availability in physiology and disease.

## Introduction

Glutathione (GSH) is the predominant antioxidant in most aerobic species.^1,2^ In mammals, it is synthesized in the cytosol in a two-step process involving glutamate and cysteine ligation followed by condensation with glycine. In addition to its antioxidant function, GSH also plays roles in disulfide bond formation, protein regulation, and iron-sulfur cluster biosynthesis, among others.^3–6^ GSH synthesis is essential for mammalian embryonic development^4,7^. While GSH depletion is associated with diseases such as type II diabetes, non-alcoholic fatty liver disease, and Parkinson’s disease, GSH levels are frequently upregulated in cancer^4,8–14^ However, whether GSH dysregulation is causative or symptomatic of disease progression in these diverse settings remains poorly understood.

Current models of GSH metabolism in disease are limited by the complex regulation of GSH abundance at the cellular level. The rate limiting enzyme, glutamate-cysteine ligase, is subject to feedback inhibition by GSH itself as well as cysteine availability.^5,15,16^ Additionally, both steps of GSH synthesis are regulated transcriptionally and post-transcriptionally.^14^ While genetic and pharmacological inhibition of GSH synthesis has provided insight into the consequences of GSH depletion, it has been very difficult to restore GSH abundance in disease models.^17,18^ GSH is not readily imported into cells, and is rapidly cleared from the serum.^19–21^ As a result, methods to increase GSH availability in disease models have been limited to global provision of cell-permeable GSH esters or GSH precursors, which require functional GSH synthesis and can act as antioxidants on their own.^4,22,23^ Additionally, activation of the Nrf2-Keap1 pathway to upregulate the expression of enzymes in the GSH synthesis pathway also impacts other downstream effectors involved in the cellular antioxidant response.^14,24,25^ Finally, none of these methods can discern whether specific subcellular GSH pools have differential roles in disease progression.

In recent years, engineered bacterial enzymes such as NDI1 and *Lb*NOX have increasingly been used to interrogate the role of oxidative metabolism in complex biological systems.^26,27^ Here, we demonstrate that bacterial enzymes can similarly be used to manipulate compartment-specific GSH pools in cells and mouse models.

## Results

### An engineered bacterial enzyme to study oxidative cell death

Mammalian GSH synthesis is a two-step process regulated transcriptionally, post-transcriptionally, and by feedback inhibition on the rate-limiting enzyme, glutamate-cysteine ligase (Fig. 1A). Conversely, some bacterial species express a single bifunctional enzyme, GshF, that possesses both glutamate-cysteine ligase and glutathione synthetase activities (Fig. 1A). In particular, the GshF enzyme expressed in *Streptococcus thermophilus* is not subject to feedback inhibition, allowing rapid accumulation of GSH when expressed in bacterial and mammalian systems^28,29^. To evaluate the use of this enzyme in manipulating compartment-specific GSH pools, we transduced HEK 293T cells with either untargeted or mitochondria-targeted, human codon-optimized *GshF*, henceforth referred to respectively as *GshF* and *mito-GshF*, and confirmed their subcellular localization by cell fractionation (Fig. 1B).

**Figure 1.**
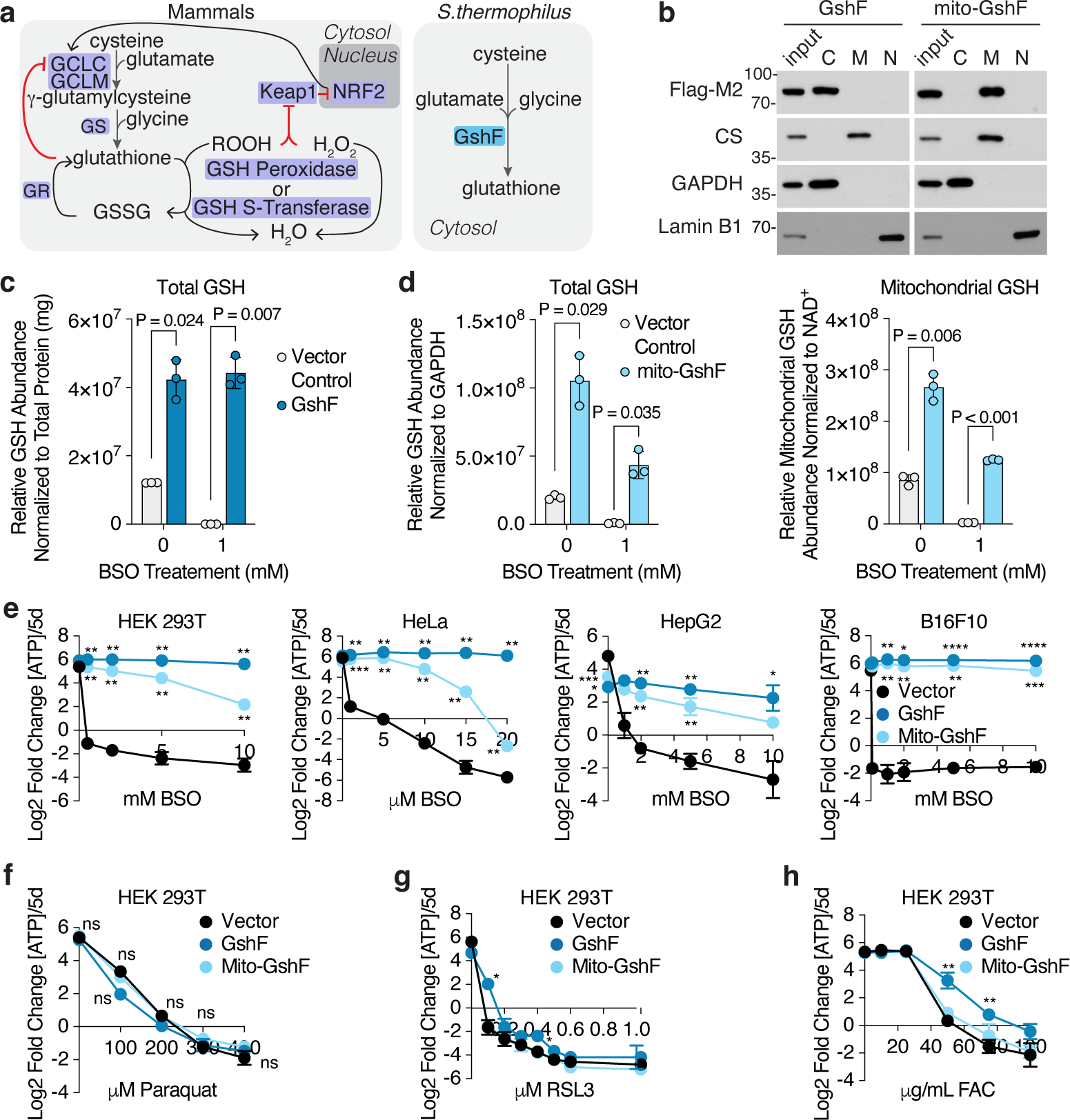
Expression of an engineered bacterial enzyme protects cells from death due to inhibition of GSH synthesis, but not ferroptosis. A. Schematic of glutathione (GSH) synthesis and regulation in mammals compared to *Streptococcus thermophilus.* GCLC, glutamate cysteine ligase catalytic subunit. GCLM, glutamate cysteine ligase modifying subunit. GR, glutathione reductase. B. The localization of engineered GshF proteins was determined by immunoblotting subcellular fractions from HEK 293T cells expressing cDNA constructs of either *GshF* or *mito-GshF*. The cytosolic, membrane-bound organelle, and nuclear and cytoskeletal fractions are denoted by C, M, and N, respectively. CS, Citrate Synthase. C. Total GSH abundance in HEK 293T cells expressing indicated cDNAs and treated with 1mM buthionine sulfoximine (BSO) for 48 hours, normalized by total protein. D. Total (left) and mitochondrial (right) GSH abundance in HEK 293T cells expressing indicated cDNAs and treated with 1mM BSO for 48 hours, normalized by GAPDH levels (total) or NAD^+^ levels (mitochondria). E. Fold change in total ATP (log_2_) of indicated cell lines expressing a vector control or specified cDNAs and treated for 5 days with the denoted concentrations of buthionine sulfoximine (BSO). Fold change in ATP is a proxy for cell doublings. P < 0.05 are indicated by asterisks according to GraphPad style. F. Fold change in total ATP (log_2_) of HEK 293T cells expressing a vector control or indicated cDNAs and treated for 5 days with the indicated concentration of paraquat. Fold change in ATP is a proxy for cell doublings. G. Fold change in total ATP (log_2_) of HEK 293T cells expressing a vector control or indicated cDNAs and treated for 5 days with the indicated concentration of RSL3. Fold change in ATP is a proxy for cell doublings. P < 0.05 are indicated by asterisks according to GraphPad style. H. Fold change in total ATP (log_2_) of HEK 293T cells expressing a vector control or specified cDNAs and treated for 5 days with the indicated concentrations of ferric ammonium citrate (FAC). Fold change in ATP is a proxy for cell doublings. P < 0.05 are indicated by asterisks according to GraphPad style.

To determine the efficacy of GshF-mediated GSH synthesis, we quantified GSH abundance from HEK 293T cells expressing either *GshF* or *mito-GshF* in the presence or absence of the mammalian GSH synthesis inhibitor buthionine sulfoximine (BSO) (Fig. S1A). Expression of either *GshF* or *mito*-*GshF* increased GSH abundance in cells grown in standard medium and maintained cellular GSH levels in the presence of BSO (Fig. 1C, 1D, S1B). *Mito-GshF* expression was also sufficient to rescue mitochondrial GSH abundance after treatment with BSO (Fig. 1D). Both *GshF*- and *mito-GshF* expression prevented BSO-induced cell death up to millimolar amounts of drug treatment in a panel of human and mouse cell lines (Fig. 1E).

GSH depletion can induce ferroptosis, a programmed cell death pathway caused by accumulation of lipid peroxides.^30–33^ The central line of defense against lipid peroxide accumulation is GPX4, which converts lipid peroxides to lipid alcohols at the expense of GSH (Fig. S1A).^33^ We asked whether GSH availability is limiting during ferroptosis in mammalian cells.^34^ To address this, we treated *GshF* and *mito-GshF* expressing cells with oxidizers and ferroptosis inducers (Fig. S1A). Surprisingly, *GshF* and *mito-GshF* expression failed to protect cells from hydrogen peroxide and paraquat exposure, which both induce reactive oxygen species (ROS) (Fig. 1F, S1A, S1C). *GshF* and *mito-GshF*-expressing cells also remained sensitive to RSL3-mediated inhibition of GPX4, which repairs lipid peroxide to prevent ferroptosis (Fig. S1A, 1G). However, *GshF* expression protected cells from iron overload induced by ferric ammonium chloride (FAC) (Fig. 1H, S1A). These data suggest that increased GSH availability is not sufficient to prevent ferroptosis and is not a limiting factor in this process.

### Targeted CRISPR-Cas9 screens under compartment-specific GSH limitation

To investigate compartment-specific vulnerabilities in cells with uncoupled mitochondrial and cytosolic GSH pools, we performed a metabolism-targeted CRISPR-Cas9 screen in Jurkat cells expressing *GshF* or *mito-GshF* in which the recently identified mitochondrial GSH transporter, *SLC25A39,* was knocked out (Fig. 2A, Fig. S2A, S2B).^28^ Cells were grown in standard RPMI or treated with a dose of BSO lethal to *SLC25A39*-knockout cells complemented with a guide-resistant *SLC25A39* cDNA (Fig. 2B). Both *GshF*- and *mito-GshF-* expressing cells were sensitive to depletion of genes involved in ferroptosis and iron metabolism, confirming that GSH is not limiting for ferroptosis (Fig. 2C). Additionally, neither *GshF* nor *mito-GshF* expression was sufficient to rescue superoxide dismutase (*Sod1*) depletion or defects in iron sulfur cluster biosynthesis resulting from cysteine desulfurase (*Nfs1*) knockout (Fig. 2C). As expected, *SLC25A40,* another recently identified mitochondrial GSH transporter, was the top hit in *GshF-*expressing *SLC25A39*-knockout cells compared to those expressing *mito-GshF* irrespective of BSO treatment (Fig. 2C, S2C).^28^ Under BSO treatment, genes related to lipid metabolism and ferroptosis were essential in *GshF-*expressing cells, which may indicate that having lower mitochondrial GSH levels may sensitize cells to lipid peroxides in the mitochondria (Fig. S2D). Conversely, the genes most essential in cells with high mitochondrial GSH but with low cytosolic GSH due to BSO treatment were largely related to oxidative phosphorylation, indicating that cells with high mitochondrial GSH are still sensitive to mitochondrial dysfunction (Fig. S2E). Together, these data indicate that cytosolic and mitochondrial GSH fulfill distinct essential roles.

**Figure 2.**
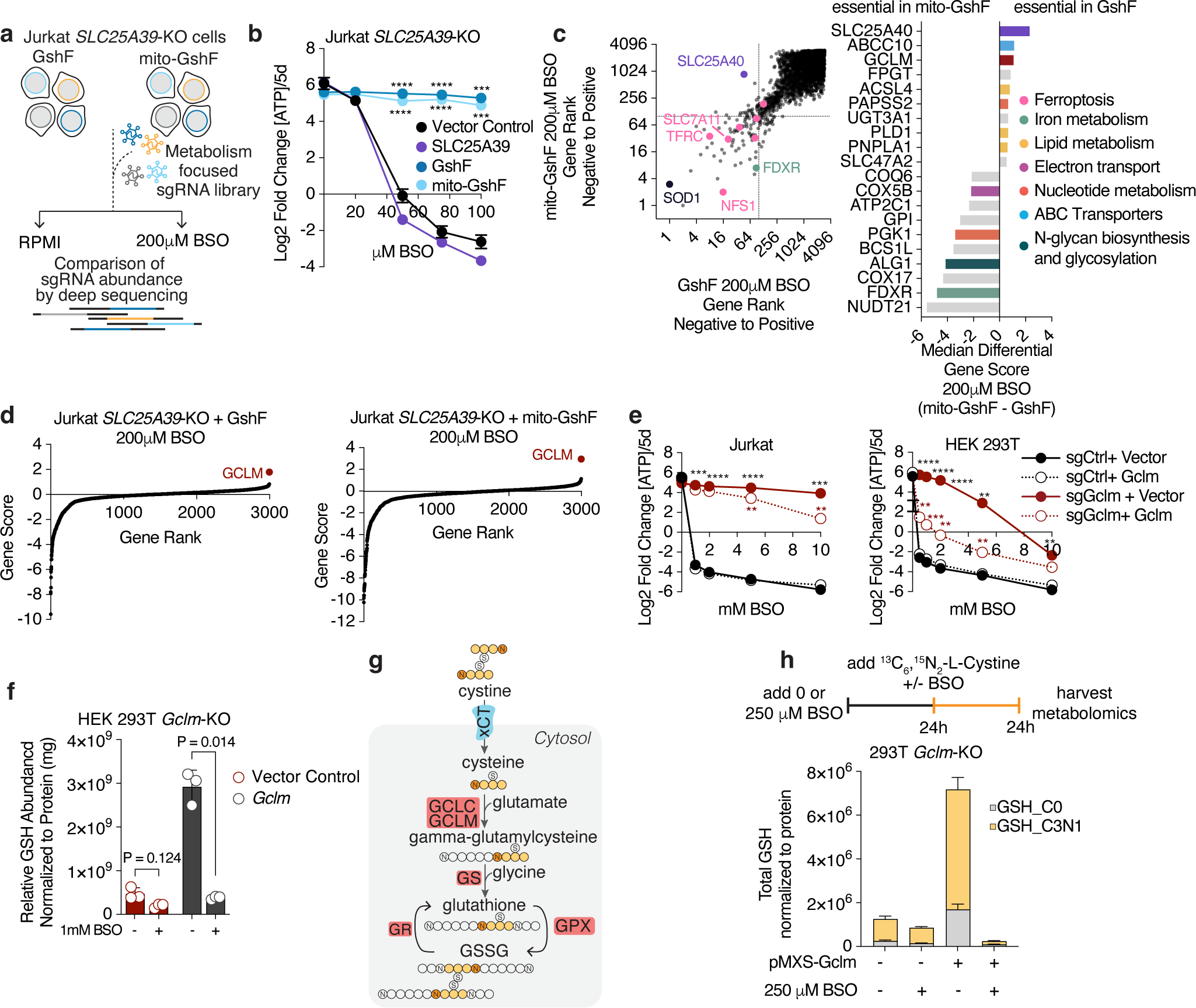
A metabolism-focused CRISPR-Cas9 screen identifies differential roles of subcellular GSH pools and mechanisms of BSO resistance. A. Schematic of metabolism-targeted CRISPR-Cas9 screen in Jurkat *SLC25A39*-knockout cells. B. Fold change in total ATP (log_2_) of Jurkat *SLC25A39*-knockout cells expressing a vector control or indicated cDNAs and treated for 5 days with the indicated concentrations of BSO. Fold change in ATP is a proxy for cell doublings. P < 0.05 are indicated by asterisks according to GraphPad style. C. Left, plot of gene score ranks from Jurkat *SLC25A39*-knockout screens treated with 200μM BSO, where genes are ranked from those with the most enriched sgRNAs to the most depleted. Genes essential in *SLC25A39*-knockout cells expressing both *GshF* and *mito-GshF* are in the lower left quartile, those specifically essential in *GshF*-expressing cells are in the top left quartile, and those specifically essential in *mito-GshF*-expressing cells are in the bottom right quartile. Right, top 10 genes scoring as differentially required upon BSO treatment in cells expressing *GshF* compared to *mito-GshF*, colored by association with indicated pathways. D. Gene scores of Jurkat *SLC25A39-*knockout cells expressing *GshF* (left) or *mito-GshF* (right) and treated with 200μM BSO for 14 doublings. The gene score is calculated as the median log_2_ fold change in all sgRNAs targeting a specific gene during the course of the culture period, as compared to a sample taken immediately before treatment began. E. Relative fold change in the total ATP (log_2_) of Jurkat (left) and HEK 293T cells (right) transduced with indicated sgRNAs and either complemented with sgRNA-resistant *Gclm* cDNA or an empty vector control. Cells were treated for 5 days with the indicated BSO doses. P < 0.05 are indicated by asterisks according to GraphPad style; comparisons of sg*Gclm* + Vector Control vs. *Gclm* cDNA are colored red and comparisons of sg*Gclm* vs. sg*Ctrl* are colored black. F. Left, relative abundance of GSH in *Gclm*-knockout HEK 293T cells complemented with sgRNA-resistant *Gclm* cDNA or an empty vector control and treated with 1mM BSO for the indicated times. Data is normalized to the total abundance of GSH in each cell line treated with BSO for 0h. Right, GSH/GSSG ratio at each timepoint of BSO treatment in indicated cell lines. G. Schematic describing incorporation of ^13^C_6_,^15^N_2_-L-Cystine into glutathione. H. Top, schematic of ^13^C_6_,^15^N_2_-L-Cystine tracing experiment. Cells were treated with 0μM or 250μM BSO for 24 hours prior to ^13^C_6_,^15^N_2_-L-Cysteine addition. Cells were maintained with 0μM or 250μM BSO for an additional 24h in the presence of ^13^C_6_,^15^N_2_-L-Cystine before quantification of metabolites. Bottom, total incorporation of cystine into GSH in the presence or absence of 250μM BSO.

Surprisingly, loss of the non-essential subunit of glutamate-cysteine ligase (GCL), *Gclm,* enabled both cell lines to proliferate in the presence of BSO (Fig 2D). Gclm lowers the K_m_ of GCL for glutamate and increases the K_i_ of GSH but is not required for catalytic function.^14^ To determine whether loss of *Gclm* could be a generalizable mechanism of BSO resistance, we generated *Gclm*-knockouts in wild-type HEK 293T and Jurkat cells. Indeed, loss of *Gclm* conferred resistance to BSO, and complementation of *Gclm*-knockout cells with a guide-resistant *Gclm* cDNA attenuated BSO resistance (Fig. 2E, Fig. S3A). We performed metabolomics analysis on *Gclm-*knockout cells treated with BSO and confirmed that *Gclm*-knockouts, but not those complemented with *Gclm* cDNA, maintained their GSH abundance in the presence of BSO (Fig. 2F). To determine whether *Gclm-* knockout cells continued to synthesize glutathione in the presence of BSO, we measured cystine incorporation into GSH after pre-treating cells with BSO for 24 hours (Fig 2G). *Gclm*-knockout cells, but not those complemented with *Gclm* cDNA, were able to incorporate ^15^N_2_,^13^C_6_-L-Cystine into glutathione even after pre-treatment with BSO for 24 hours (Fig 2H, S3B). Altogether, these data show that loss of *Gclm* enables resistance to BSO.

### Increased GSH availability is incompatible with early mouse development

Increasing glutathione pools *in vivo* has been challenging due to the complex regulation of GSH synthesis. GSH is not orally available, and supplemented GSH is cleared from the bloodstream within minutes.^21,35,36^ Additionally, genetic methods to increase GSH are limited to manipulation of multifaceted antioxidant response pathways, further complicating interrogation of GSH sufficiency.^14,24,25^ To determine whether *GshF* expression could increase GSH abundance *in vivo*, we generated genetically engineered mice by inserting either *GshF* or *mito-GshF* under control of a STOP codon flanked by two *LoxP* sites at the Rosa26 locus (Fig. 3A, S4A). To increase GSH availability in all tissues, we mated heterozygous floxed mice to a CMV-Cre model. CMV-mito-GshF^Δ/0^ pups were born at expected ratios and appeared healthy but did not exhibit increased GSH concentration in any tissues despite expressing *mito-GshF* (Fig. S4B-S4D). Conversely, CMV-GshF^Δ/0^ mice were greatly underrepresented at P0, and the few CMV-GshF^Δ/0^ pups born on P0 died within 12 hours of birth (Fig. 3B, S5A). We assessed the presence of CMV-GshF^Δ/0^ embryos at several stages and found a general trend of decreasing numbers of CMV-GshF^Δ/0^ embryos as development proceeded (Fig. 3B). We examined CMV-GshF^Δ/0^ and their CMV-GshF^0/0^ littermates at E17.5 and found that CMV-GshF^Δ/0^ embryos were slightly shorter than their CMV-GshF^0/0^ littermates but did not observe histologic differences in any of the tissues (Fig. S5B-D). These results indicate that GSH homeostasis is essential in early development and that increased GSH levels during embryogenesis are not compatible with life.

**Figure 3.**
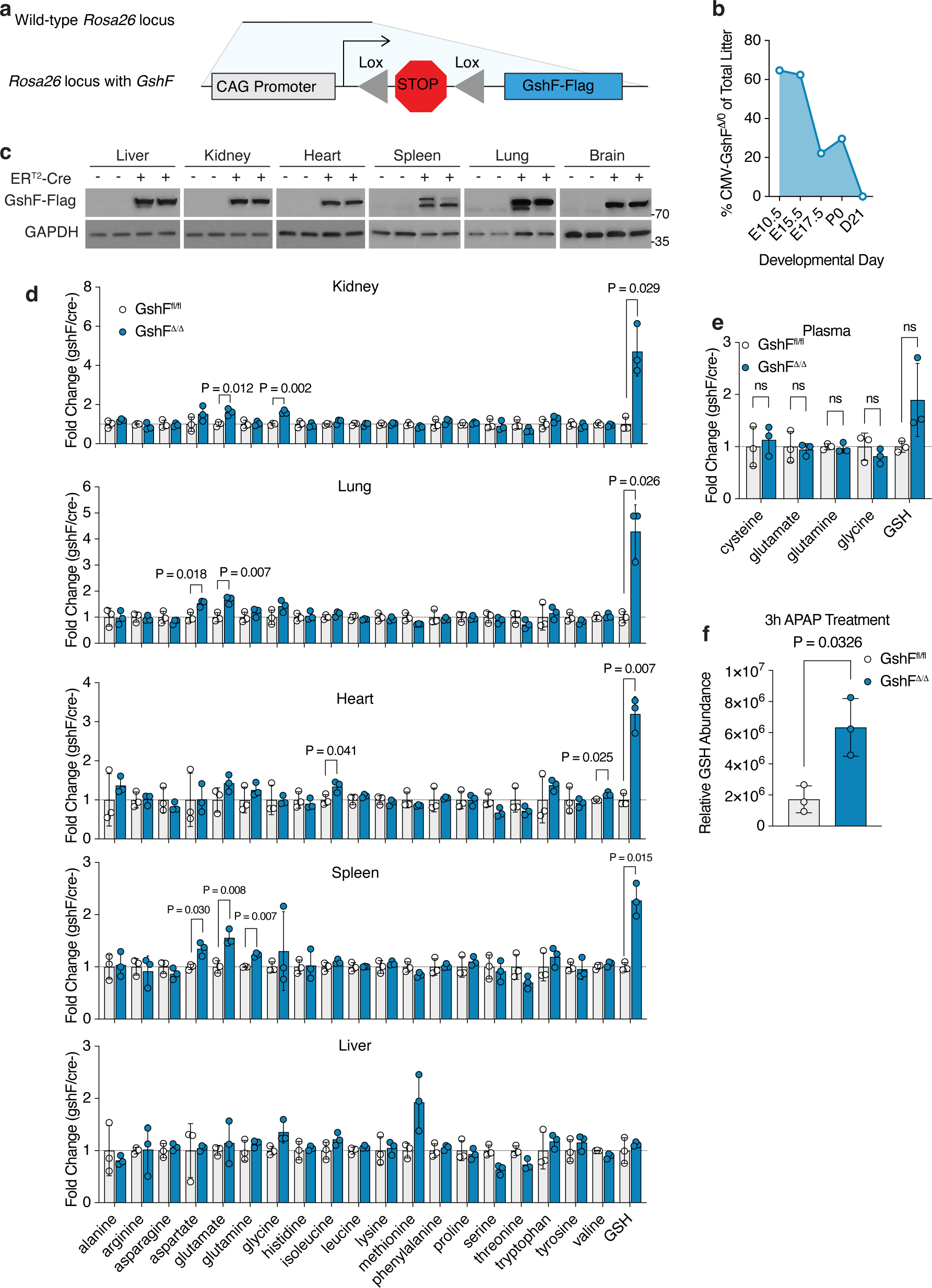
Generation of a mouse with constitutively high glutathione. A. Schematic of *GshF* insertion into the Rosa26 locus of CB57bl/6 mice. B. Percent of CMV-GshF^Δ/0^ of the total litter at indicated developmental stages. Mendelian genetics predict 50%. C. Immunoblots of indicated tissues from littermates treated with 75mg/kg of tamoxifen for 5 consecutive days. D. Abundance of indicated metabolites in each tissue from male ER^T2^-GshF^Δ/Δ^ or GshF^fl/fl^ littermates treated with 75mg/kg of tamoxifen for 5 consecutive days. For clarity, only P values < 0.05 are displayed on the charts. E. Metabolomics analysis of indicated plasma metabolites from male ER^T2^-GshF^Δ/Δ^ or GshF^fl/fl^ littermates before and after tamoxifen induction. F. GSH abundance in livers from ER^T2^-GshF^Δ/0^ or ER^T2^-GshF^0/0^ mice treated with acetaminophen (APAP) for 3 hours.

### Mice with constitutively high glutathione levels are viable

To examine the impact of GSH accumulation in adult mice, we mated GshF^fl/0^ mice to a tamoxifen-inducible ER^T2^-Cre model and induced GshF expression once the mice were four weeks old. The body and tissue weights of ER^T2^-GshF^Δ/0^ mice were comparable to their ER^T2^-GshF^fl/0^ littermates, and we did not observe any signs of distress or death for four months (Fig. S6A, S6B). We confirmed that GshF was indeed expressed in the tissues of ER^T2^-GshF^Δ/0^ mice by Western blot (Fig. 3C). To determine the effect of GshF expression on tissue metabolism, we analyzed plasma and tissue metabolites from ER^T2^-GshF^Δ/Δ^ and their GshF^fl/fl^ littermates and found that in every tissue except the liver and plasma, GSH was more abundant in mice expressing *GshF* (Fig. 3D, 3E, S6C). The extent to which GSH was increased in each tissue varied, with the kidney and the lung exhibiting the greatest increase in GSH abundance compared to those from *GshF*-null mice. Importantly, the abundance of amino acids in *GshF*-expressing tissues was largely comparable to those in *GshF*-null tissue, including the precursors of GSH synthesis, glutamate and glycine (Fig. 3D). In humans, liver GSH is rapidly depleted in acetaminophen (APAP) overdose, and provision of the GSH precursor N-acetylcysteine is the current standard of care.^37^ To determine whether GshF was active in the liver, we injected mice with 300mg/kg APAP and measured metabolites from GshF-expressing and GshF-null littermates.^37,38^ GshF-expressing mice had about three times as much GSH in their liver 3h after APAP treatment (Fig. 3F). These results indicate that increased GSH availability is compatible with life and opens up the potential to examine GSH sufficiency in a variety of contexts.

Initially, we characterized the effects of high GSH on cell populations where GSH synthesis is required for cell development or activity, such as blood cells and T cells. GSH is required for iron-sulfur cluster biosynthesis and defects in GSH synthesis or GSH import into mitochondria can cause anemia.^6^ To determine whether high GSH could impact blood cell development, we performed complete blood counts on *GshF*-expressing mice. There were very few differences between the *GshF*-expressing mice and their *GshF*-null littermates, with the only significant difference being in the mean corpuscular hemoglobin concentration (MCHC), a measure of hemoglobin relative to the size of the cell (Fig. S6D). GSH synthesis is also required for T cell activation.^39,40^ To determine whether *GshF* expression impacted immune cell development, we analyzed the T cell populations in the spleen, thymus, lungs, mesenteric lymph nodes, and colon of ER^T2^-GshF^Δ/Δ^ and GshF^fl/fl^ littermates (Fig. S7A-S7J). There were no significant differences in any of the immune populations between ER^T2^-GshF^Δ/Δ^ and GshF^fl/fl^ mice, although we observed several non-significant trends in the colon, particularly in the lamina propria (Fig. S7A-S7J). The only trend we observed across tissues was a slight increase in TCRγδ cells in ER^T^^2^-GshF^Δ/Δ^ mice (Fig. S7J).

To unbiasedly investigate the effect of high GSH on tissue homeostasis, we next performed RNAseq analysis on the tissues with the greatest increase in GSH with *GshF* expression, the kidney and lung (Fig. 3D, 4A, B). Gene ontology (GO) analysis revealed that the only common pathway differentially expressed in both the kidney and the lung was response to steroid hormones, which was downregulated in both tissues (Fig. 4A-D). Several significant GO terms in lung were related to an increased immune response and to an increase in genes contributing to the extracellular matrix (Fig. 4A, B). In the kidney, differential gene expression corresponded with cell differentiation and lipid metabolism, as well as membrane transport (Fig. 4C, D). Some of the downregulated lipid metabolism genes are involved in steroid biosynthesis, which may contribute to the downregulation of the steroid hormone response in both tissues. Future work will need to determine whether GSH has differential functions in individual tissues.

**Figure 4.**
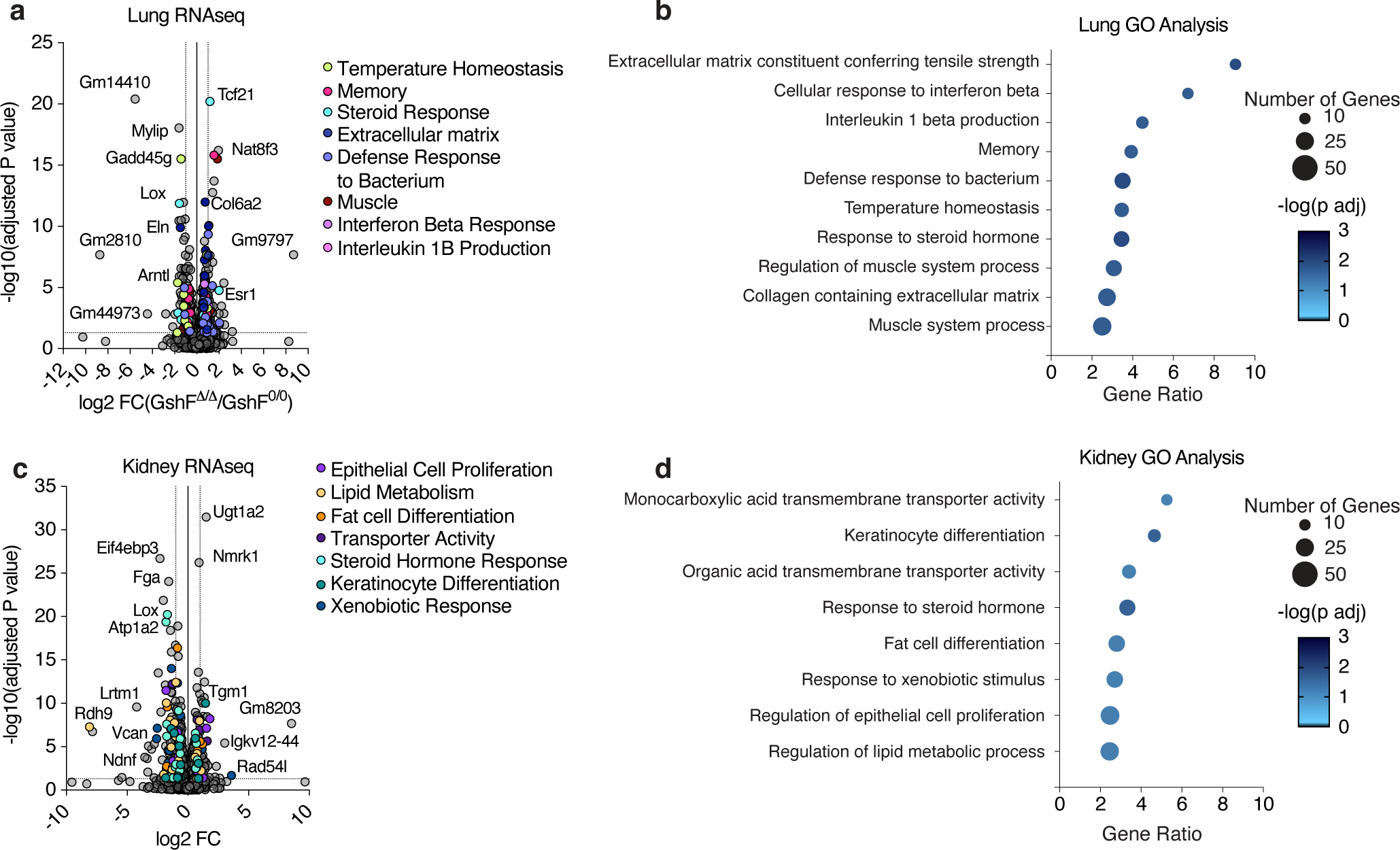
RNAseq of GshF mice reveals common and tissue-specific pathways impacted by high GSH. A. RNAseq of lung tissues from male ER^T2^-GshF^Δ/Δ^ vs. GshF^fl/fl^ littermates two weeks after completion of tamoxifen injections. B. GO analysis of top differentially expressed genes in lung. C. RNAseq of kidney tissues from male ER^T2^-GshF^Δ/Δ^ vs. GshF^fl/fl^ littermates two weeks after completion of tamoxifen injections. D. GO analysis of top differentially expressed genes in kidney.

## Discussion

Glutathione is an evolutionarily ancient antioxidant that arose around the same time as photosynthetic cyanobacteria.^41–43^ Millions of years of evolution have extended GSH’s role as a central reducing agent, and systems developed coupling GSH to critical cell processes including iron homeostasis, electron transport, and reducing power in the form of NADPH.^43–45^ Interestingly, GSH dysfunction is associated with a wide range of pathologies. GSH depletion is observed in neurodegenerative diseases, metabolic diseases like type II diabetes, chronic immunodeficiencies like AIDS, anemia, liver diseases, and lung disease, among others.^3,46,47^ Similarly, GSH abundance increases in many cancers and correlates with disease progression and mortality.^8–10,48–50^ Given the strong association of GSH dysfunction with disease and the diversity of pathological presentation, determining the critical processes impacted by changes in GSH availability remains crucial.

Genetically encoded tools to manipulate cellular metabolites *in vivo* have been useful in determining the role of specific molecules, particularly in oxidative metabolism.^26,27,51^ We have described here the design and validation of GshF to manipulate compartmentalized GSH concentration. Previous work from our and other labs has demonstrated that mitochondrial GSH is critical for cell viability, and indeed we find that cells with uncoupled mitochondrial and cytosolic GSH remain sensitive to inhibition of mitochondrial GSH transport or disruption of mitochondrial functionality.^28,52^ Surprisingly, increasing total GSH did not protect cells from ferroptosis. We interpret this to mean that GSH abundance is not limiting for GPX4 activity. The exception seems to be in iron overload, where increased cytosolic GSH is protective. While this could be related to GSH-mediated repair systems, it is also possible that GSH actively complexes free iron to prevent the Fenton reaction wherein hydrogen peroxide is dismutated to more reactive oxygen radicals that can damage proteins, lipids, and nucleic acids.^53–55^ Further work will need to be done to determine the precise mechanism.

GSH availability is tightly controlled though transcriptional, post-transcriptional, and post-translational mechanisms. While GSH depletion is detrimental for cell and organismal viability, excess GSH can also cause reductive stress and lethally disrupt iron metabolism.^6,7,56^ Our work expands upon this knowledge in demonstrating that, like in yeast, there is a toxic level of GSH incompatible with mammalian development, emphasizing that homeostatic mechanisms to limit GSH production are critical for embryonic viability. Perinatal lethality can be caused by a myriad of deficiencies, and determining the precise cause of death in these mice will require many tissue-specific models. However, this phenotype indicates that unregulated GSH synthesis or high levels of GSH may disrupt certain critical aspects of development.

In the course of this work, we also identified a possible mechanism of BSO resistance. BSO is a glutamate analog originally described by Alton Meister to inhibit the rate limiting step of GSH synthesis, glutamate-cysteine ligase (GCL).^57^ In the past, several clinical trials have queried the efficacy of using BSO to sensitize tumor cells to oxidative stress to little success^58,59^ GCL is a homodimer comprised of a catalytic subunit, GCLC, and a modifying subunit GCLM that is dispensable for GSH synthesis.^60^ Our work indicates that BSO may only inhibit the heterodimeric GCLC complex, not GCLC alone. We speculate that downregulation of or loss of GCLM might enable resistance to BSO *in vivo*. Future efforts to develop GSH inhibitors specific for GCLC or may still prove clinically useful in some contexts, particularly in combination with thioredoxin inhibitors in cancer settings and in combination with anti-parasitic agents in African sleeping sickness and other *Trypanosome* diseases, which BSO has shown some promise in treating.^61,62^

Intriguingly, increased GSH availability has minimal impact on metabolism in adult animals, which provides a unique opportunity to restore GSH concentration without otherwise perturbing disease states. While there were no observable differences at baseline in ER^T2^-GshF^Δ/Δ^ mice, it is possible that there will be differential effects in specific immune challenges and disease models, although this will require a battery of tests to determine. In conclusion, our work provides a conditional mouse model that will enable dissection of the contribution of GSH sufficiency to disease progression in a variety of contexts.

## Supporting Figure Legends

**Supporting Figure 1.**
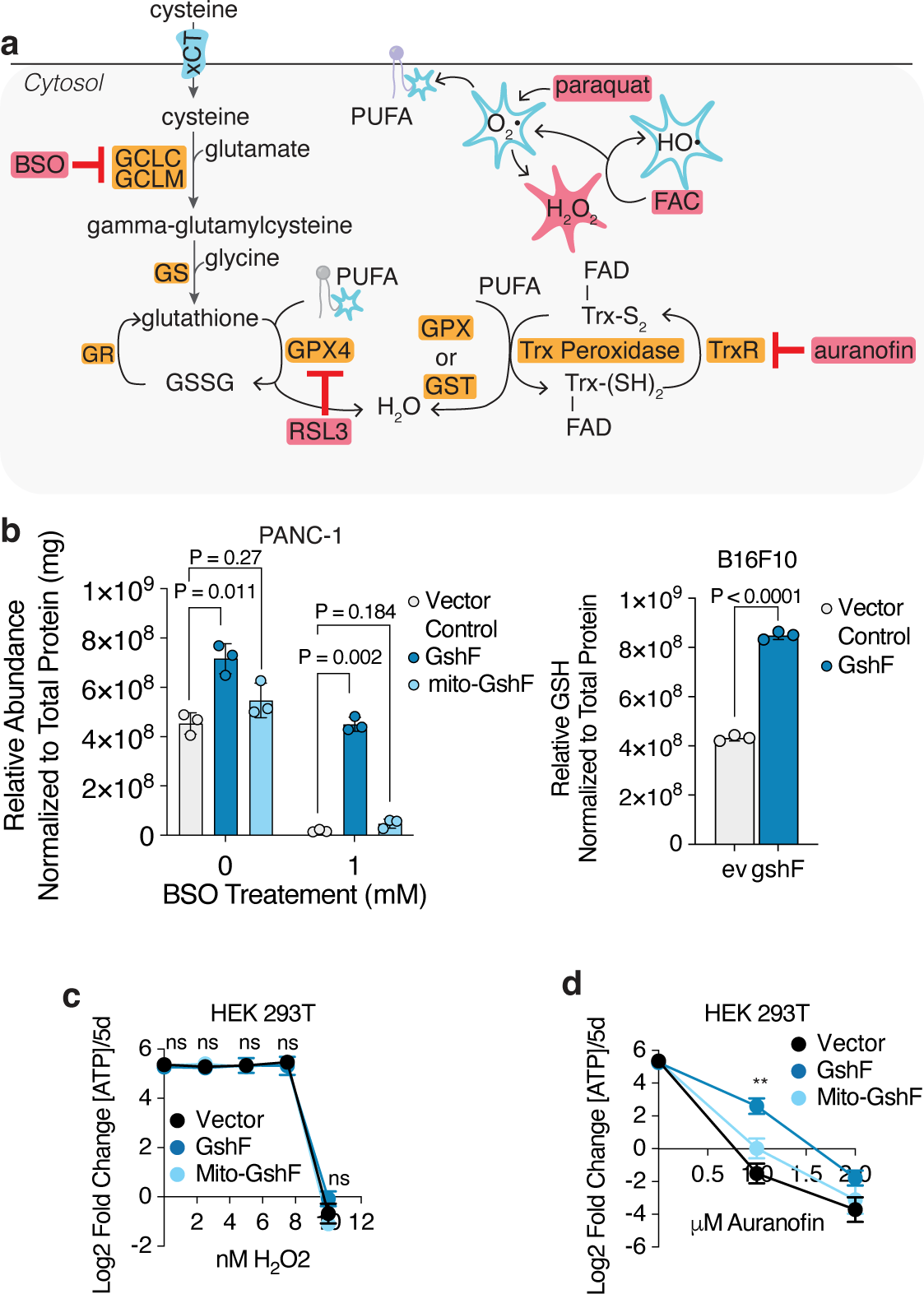
GSH availability is not limiting for the oxidative stress response. A. Schematic of enzymatic inhibitors and ROS generators impacting glutathione metabolism and ferroptosis. BSO, buthionine sulfoximine; FAC, ferric (III) ammonium citrate. B. Left, total abundance of GSH in mouse PANC-1 cells expressing indicated cDNAs and treated with 0mM or 1mM BSO for 72 hours, normalized by total protein (mg). Right, fold change in total ATP (log_2_) of PANC-1 cells expressing indicated cDNAs and treated for 5 days with the denoted concentrations of buthionine sulfoximine (BSO). Fold change in ATP is a proxy for cell doublings. P < 0.05 are indicated by asterisks according to GraphPad style. C. Total abundance of GSH in B16F10 cells expressing indicated cDNAs and treated with 0mM or 1mM BSO for 72 hours. Data is normalized by total protein (mg). D. Relative fold change in the total ATP (log_2_) of HEK 293T cells transduced with *GshF*, *mito-GshF*, or a vector control and treated for 5 days with the indicated concentrations of H_2_O_2_. E. Relative fold change in the ATP (log_2_) of indicated cells transduced with *GshF*, *mito-GshF*, or a vector control and treated for 5 days with the indicated concentrations of auranofin. P < 0.05 are indicated by asterisks according to GraphPad style.

**Supporting Figure 2.**
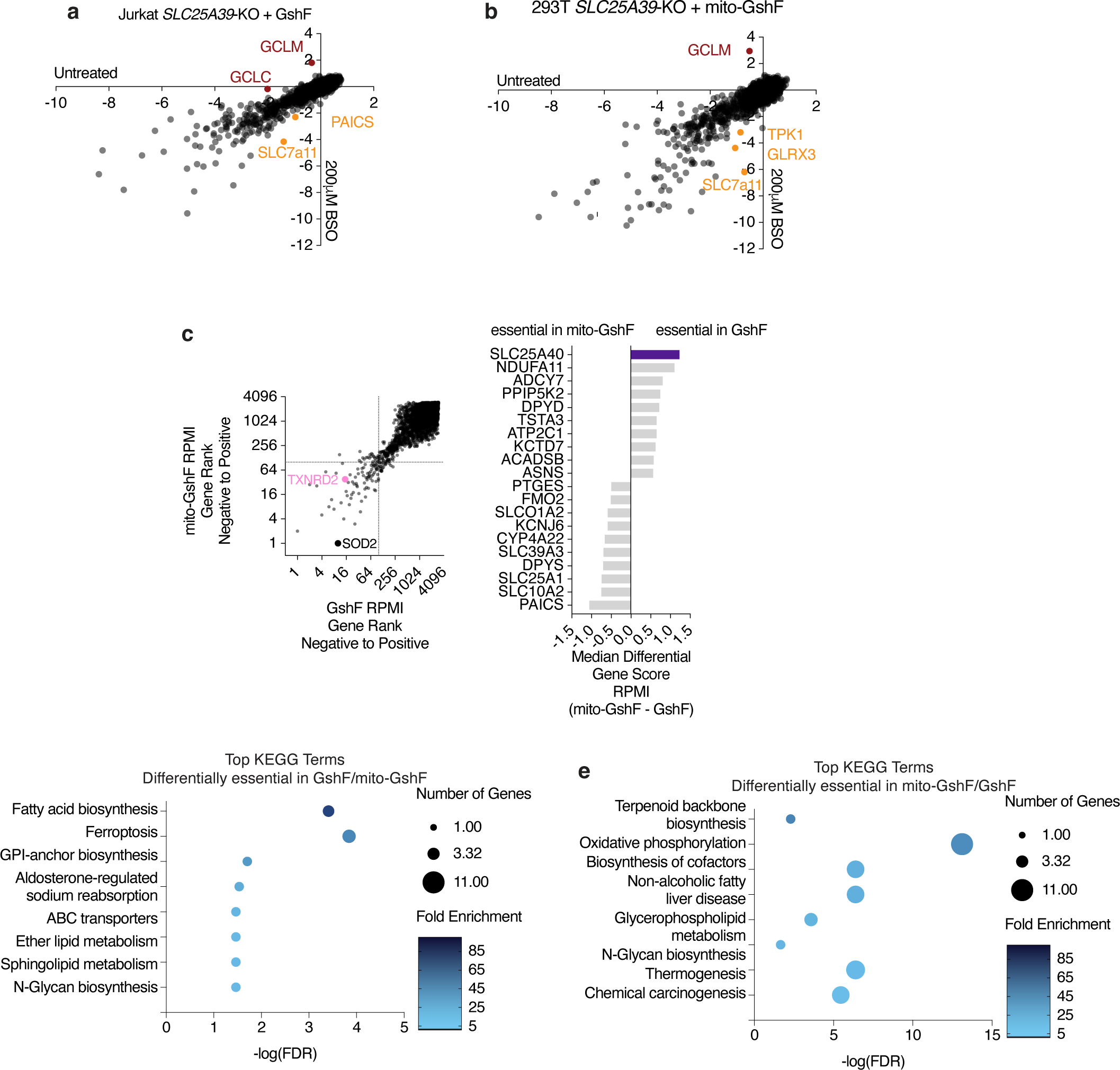
A metabolism-targeted CRISPR-Cas9 screen reveals that unregulated GSH synthesis has little impact on most metabolic pathways. A. Left, gene scores of Jurkat *SLC25A39-*knockout cells expressing *GshF* or *mito-GshF* transduced with a CRISPR-Cas9 sgRNA library targeting 3,000 metabolic genes and grown in standard media for 14 doublings. The gene score is calculated as the median log_2_ fold change in all sgRNAs targeting a specific gene during the course of the culture period, as compared to a sample taken immediately before treatment began. Right, top 10 genes scoring as differentially required in cells expressing *GshF* compared to *mito-GshF* grown in normal culture conditions. B. Gene ontology analysis of top 100 genes essential in Jurkat *SLC25A39*-knockout cells expressing *GshF* under BSO treatment. C. Gene ontology analysis of top 100 genes essential in Jurkat *SLC25A39*-knockout cells expressing *mito-GshF* under BSO treatment. D. Gene scores from Jurkat *SLC25A39*-knockout cells expressing *GshF* grown in standard medium compared to those treated with BSO for the course of the screen. E. Gene scores from Jurkat *SLC25A39*-knockout cells expressing *mito-GshF* grown in standard medium compared to those treated with BSO for the course of the screen.

**Supporting Figure 3.**
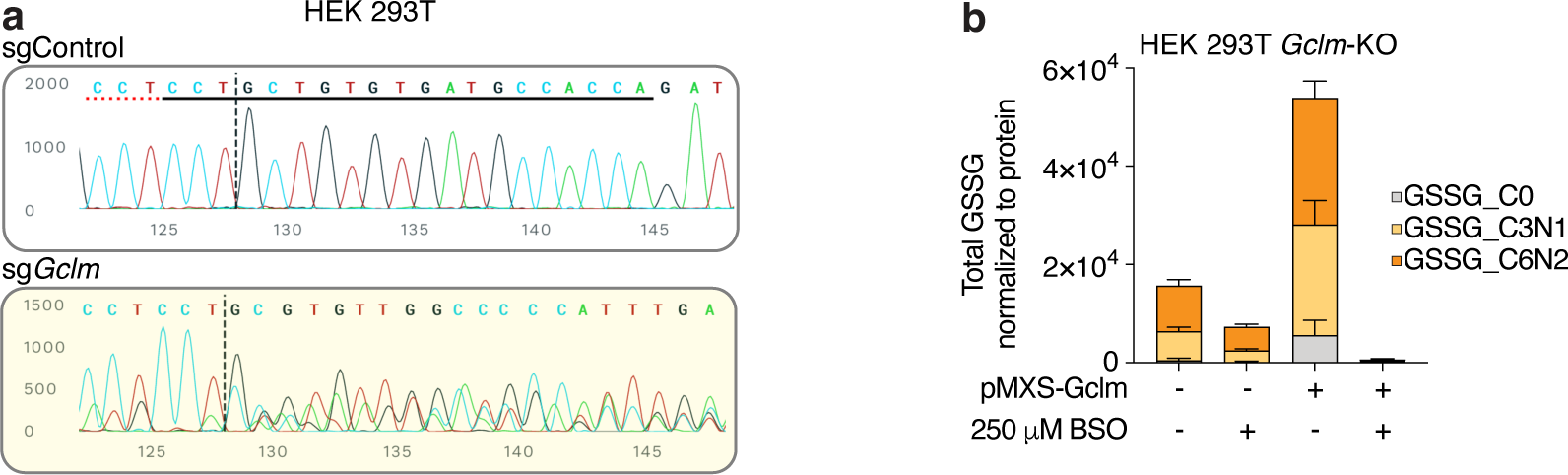
Loss of the modifying subunit of the rate-limiting enzyme in GSH synthesis provides resistance to buthionine sulfoximine. A. Sanger sequencing of genomic region targeted by sg*Gclm* in HEK 293T cells transduced with indicated sgRNAs. B. Total GSSG of indicated isotopes in the presence of absence of 250μM BSO in *Gclm*-knockout cells expressing indicated cDNAs.

**Supporting Figure 4.**
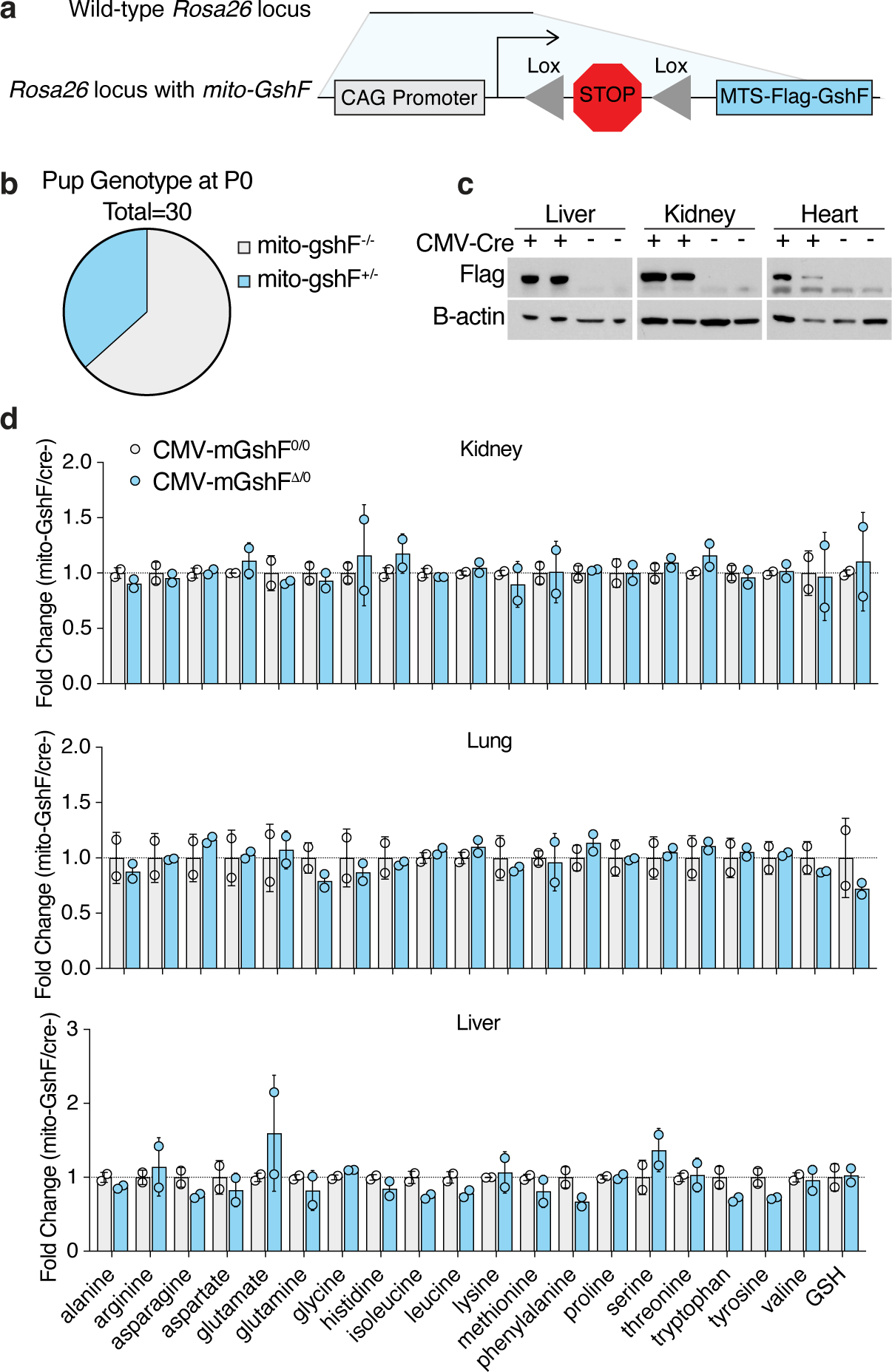
Constitutive mitochondrial glutathione synthesis is compatible with embryonic development. A. Schematic of *mito-GshF* insertion into the Rosa26 locus of CB57bl/6 mice. B. Percent of CMV-mito-GshF^Δ/0^ of the total litter. Expected ratio is 25%. C. Immunoblots of mito-GshF expression in indicated tissues from CMV-mito-GshF^Δ/0^ and CMV-mito-GshF^0/0^ littermates. D. Metabolomics analysis of indicated metabolites in CMV-mito-GshF^Δ/0^ tissues, normalized to CMV-mito-GshF^0/0^ littermate controls.

**Supporting Figure 5.**
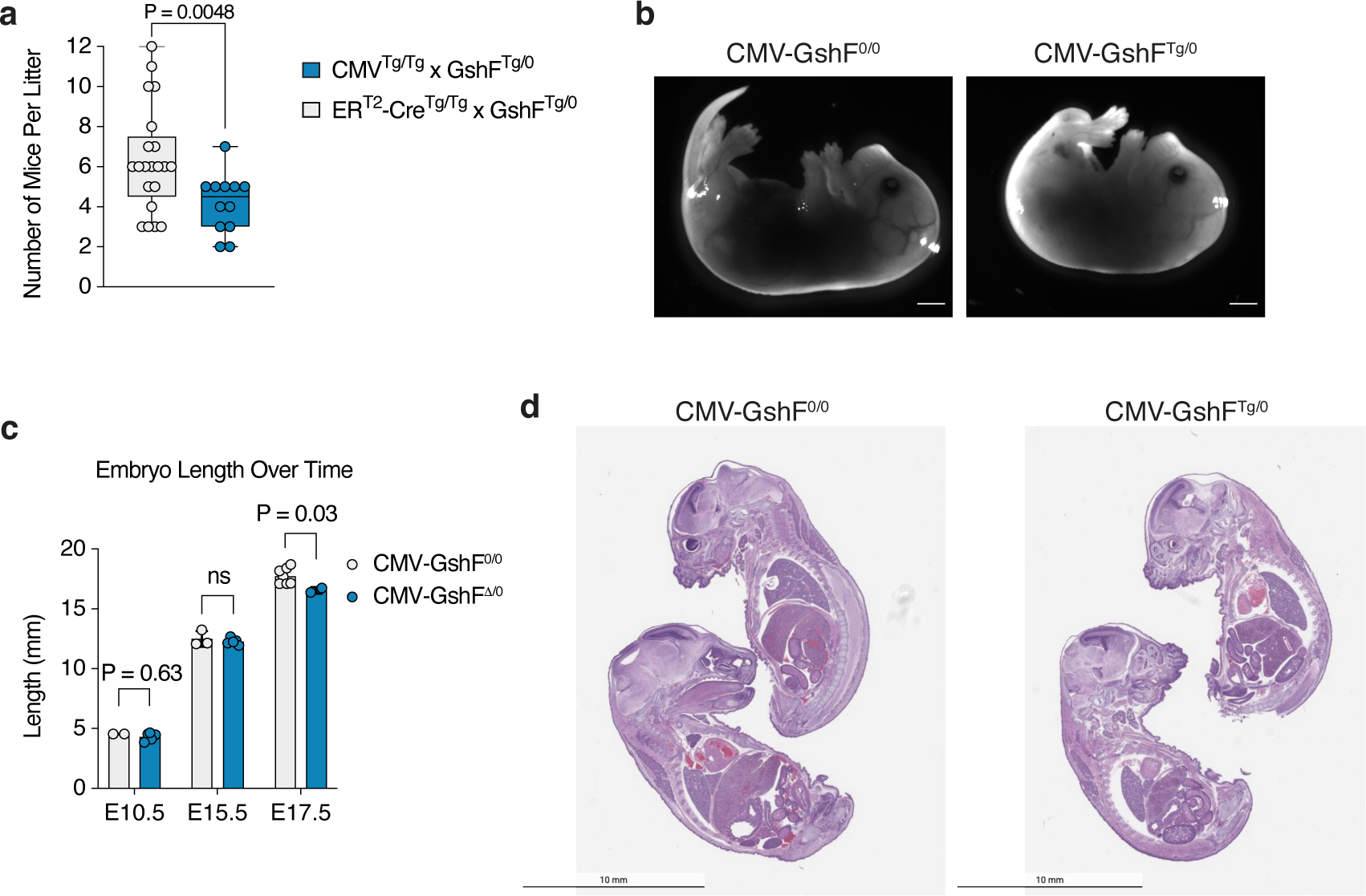
Constitutive unregulated glutathione synthesis is perinatal lethal. A. Litter size of indicated mating pairs. B. Representative image of embryos of indicated genotypes at E17.5. Scale bar is 1mm. C. Quantification of embryo length of respective genotypes at indicated stages of embryonic development. D. Representative H&E stain of E17.5 mice of indicated genotypes. Scale bar represents 10mm.

**Supporting Figure 6.**
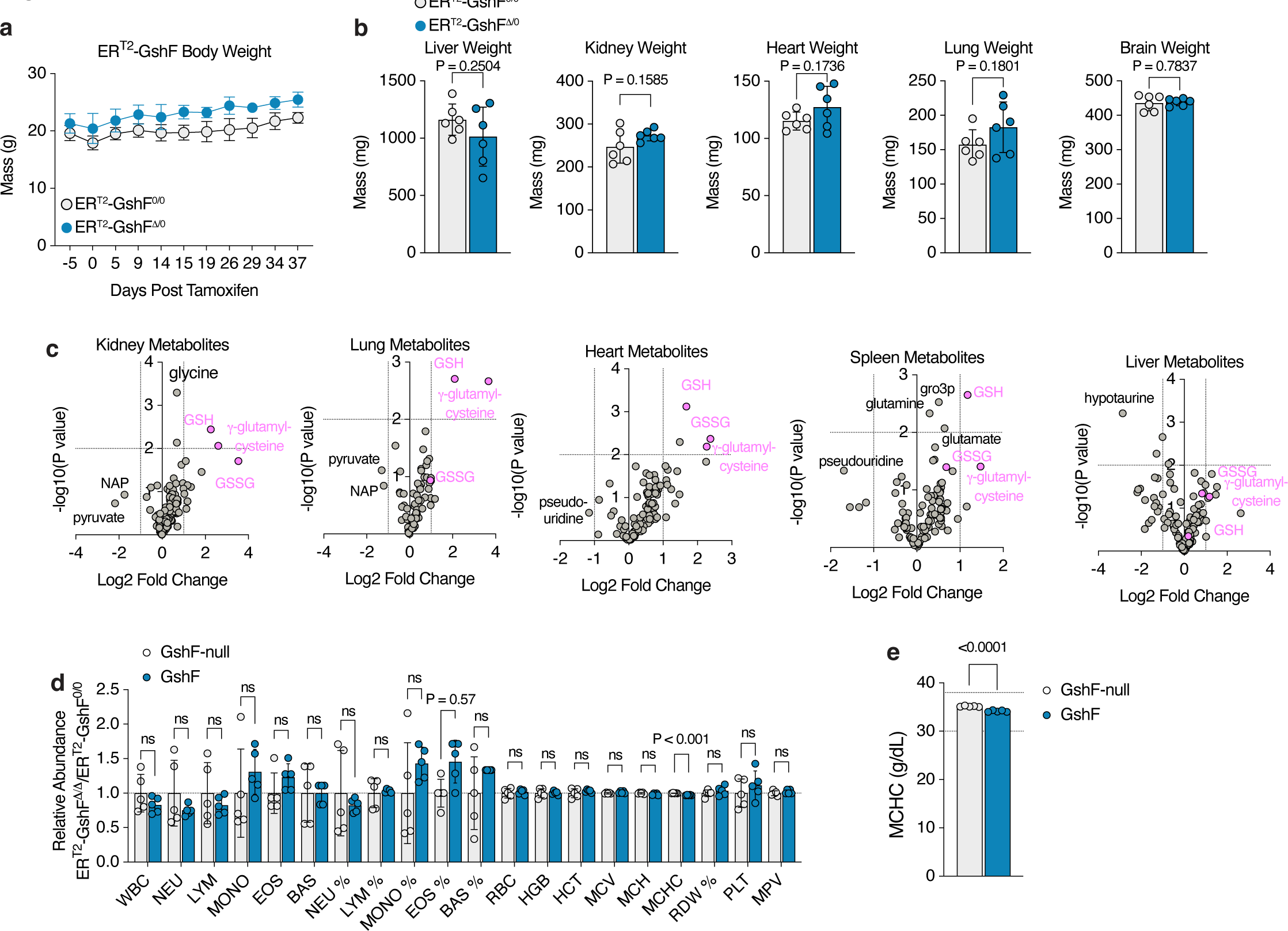
Unregulated glutathione synthesis in adult mice is compatible with life. A. Plot of body weights of littermates of indicated genotypes before and after tamoxifen treatment. B. Weights of indicated tissues two weeks after tamoxifen induction. C. Volcano plots of all metabolites detected in indicated tissues from male ER^T2^-GshF^Δ/Δ^ mice compared to GshF^fl/fl^ littermates. D. Relative blood cell types or percentages from ER^T2^-GshF^Δ/Δ^ vs. ER^T2^-GshF^0/0^ littermates. E. Relative mean corpuscular hemoglobin concentration (MCHC) cells from male ER^T2^-GshF^Δ/Δ^ vs. GshF^fl/fl^ littermates. Normal MCHC range is indicated by dotted lines.

**Supporting Figure 7.**
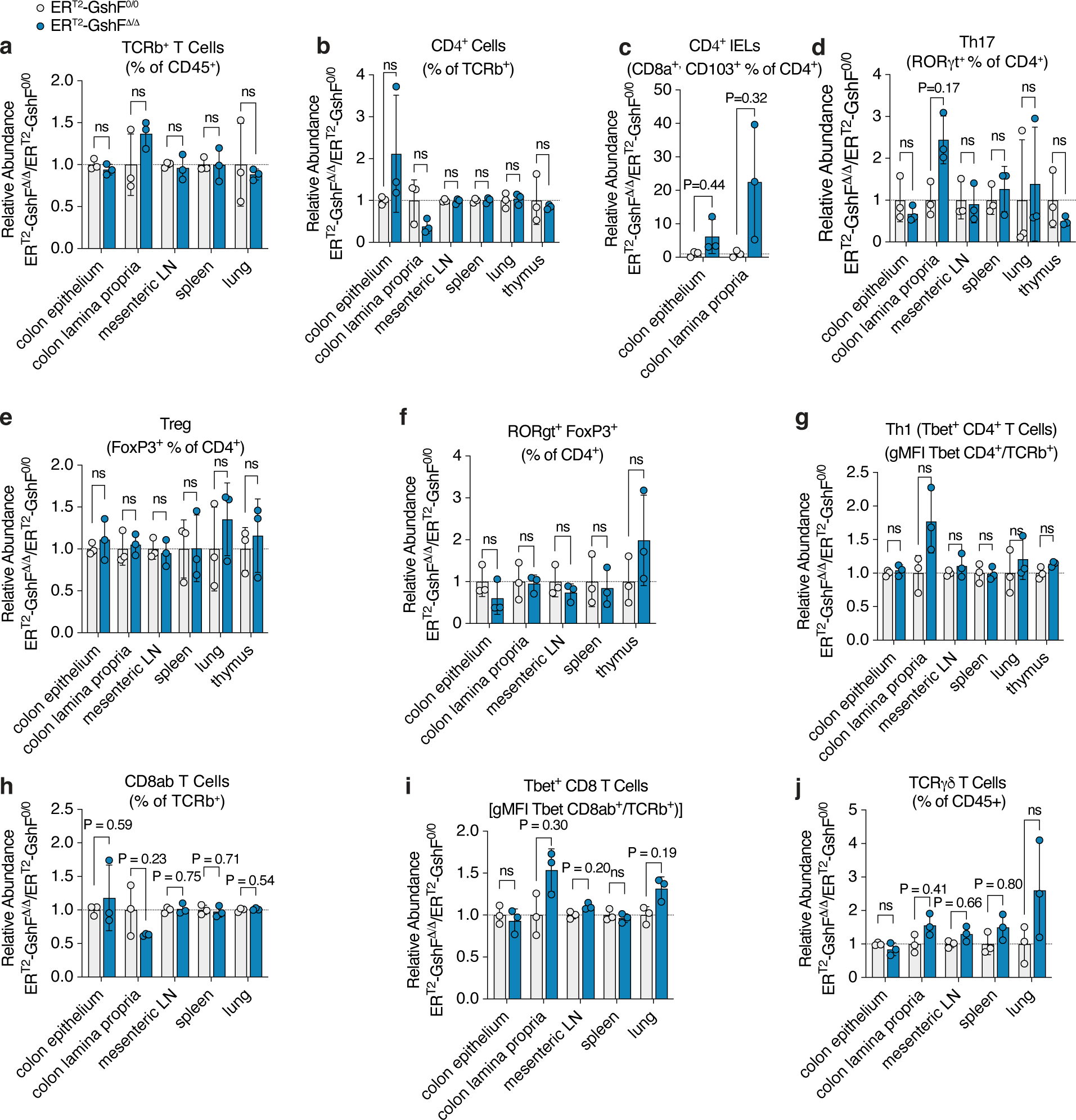
Mice with high GSH have normal immune profiles. A. Relative TCRb^+^ T cells in indicated tissues from female ER^T2^-GshF^Δ/Δ^ vs. ER^T2^-GshF^0/0^ littermates. B. Relative CD4^+^ T cells in indicated tissues from female ER^T2^-GshF^Δ/Δ^ vs. ER^T2^-GshF^0/0^ littermates. C. Relative CD4^+^ intraepithelial lymphocytes (IELs) cells in indicated colon layers from female ER^T2^-GshF^Δ/Δ^ vs. ER^T2^-GshF^0/0^ littermates. D. Relative Th17 T cells in indicated tissues from female ER^T2^-GshF^Δ/Δ^ vs. ER^T2^-GshF^0/0^ littermates. P < 0.9 are indicated numerically. E. Relative Treg cells in indicated tissues from female ER^T2^-GshF^Δ/Δ^ vs. ER^T2^-GshF^0/0^ littermates. F. Relative RORγt^+^, FoxP3^+^, CD4^+^ T cells in indicated tissues from female ER^T2^-GshF^Δ/Δ^ vs. ER^T2^-GshF^0/0^ littermates. G. Relative Th1 T cells in indicated tissues from female ER^T2^-GshF^Δ/Δ^ vs. ER^T2^-GshF^0/0^ littermates. H. Relative CD8ab^+^ T cells in indicated tissues from female ER^T2^-GshF^Δ/Δ^ vs. ER^T2^-GshF^0/0^ littermates. I. Relative CD8^+^ Tbet^+^ T cells in indicated tissues from female ER^T2^-GshF^Δ/Δ^ vs. ER^T2^-GshF^0/0^ littermates. J. Relative TCRγδ T cells in indicated tissues from female ER^T2^-GshF^Δ/Δ^ vs. ER^T2^-GshF^0/0^ littermates.

**Supporting Table S1. A metabolism-targeted CRISPR-Cas9 screen reveals that unregulated GSH synthesis has little impact on most metabolic pathways.**

All gene scores from a metabolism-targeted CRISPR-Cas9 screen performed in Jurkat *SLC25A39*-knockout cells expressing *GshF* or *mito-GshF* and treated with or without 200μM BSO.

**Supporting Table S2. Differentially expressed genes in ERT2-GshF^Δ/Δ^ vs. GshF**^0^**^/^**^0^ **kidneys.**

All base means and log_2_ fold changes of mRNAs quantified during RNAseq from ERT2^Δ/Δ^ vs. GshF^0/0^ kidneys.

**Supporting Table S3. Differentially expressed genes in ERT2-GshF^Δ/Δ^ vs. GshF**^0^**^/^**^0^ **lungs.**

All base means and log_2_ fold changes of mRNAs quantified during RNAseq from ERT2^Δ/Δ^ vs. GshF^0/0^ lungs.

## Experimental Procedures

### Cell Lines and Reagents

All cell lines were purchased from the ATCC. Cell lines were regularly monitored to be free of mycoplasma contamination and the identities of all were verified by STR profiling. All cells except HepG2 were maintained in RPMI 1640 media (Gibco) containing 2mM glutamine, 10% fetal bovine serum, 1% penicillin and streptomycin. HepG2 cells were maintained in DMEM media (Gibco) containing 4.5 g/L glucose, 110 mg/L pyruvate, 4mM glutamine, 10% fetal bovine serum, 1% penicillin and streptomycin. All cells were maintained at 37°C, 21% oxygen, and 5% carbon dioxide.

Antibodies against GAPDH (GTX627408, 1:1000) were obtained from GeneTex; FLAG M2 (F1804, 1:1000) from Sigma; Citrate synthase (14309S, 1:1000), Histone H3 (4499S), GAPDH (2118L) from Cell Signaling Technologies.

[^13^C_6_, ^15^N_2_]-L-Cystine (CNLM-4244-H-PK) and [^12^C_2_, ^15^N]-GSH (CNLM-6245-50) were purchased from Cambridge Isotope Laboratory. BSO (B2515), ferric ammonium citrate (FAC, F5879), and paraquat (856177) were purchased from Sigma Aldrich. RSL3 (S8155) and erastin (S7242) were purchased from Selleckchem. Hydrogen peroxide (1000035784) was purchased from Cumberland Swan. Auranofin (15316) and tamoxifen (13258) were purchased from Cayman Chemical Company.

### Generation of knockout cell lines

As described in Wang et al^28^, sgRNAs were cloned into BsmB1-linearized lentiCRISPR-v1-GFP (Addgene) by T4 DNA ligase (New England Biolabs). Presence of sgRNA was determined by U2 sequencing primers.

Lentiviral particles were produced by transfecting HEK 293F cells with sgRNA-containing vector, viral packaging vector deltaVPR, and lentiviral envelope vector CMV VSV-G using XtremeGene 9 transfection reagent (Roche). Viral particles were collected 48h after transfection and passed through a 0.45 μm filter to eliminate cells and cell debris. Virus was then delivered to target cell lines seeded in 6-well tissue culture plates in medium containing 8 μg/mL polybrene. Cells were centrifuged at 2200 rpm for 1hr to improve infection efficiency. Virus was removed 24 hours after transduction, and top 1% of GFP+ cells were collected by fluorescence-activated cell sorting on a BD FACSAria II. Cells were validated for loss of the target protein by immunoblotting except *Gclm*-knockout cells, which had an interfering band at the same molecular weight as Gclm. The genomic region targeted by sg*Gclm* was PCR-amplified and submitted to Sanger sequencing to ensure that the exon sequence was disrupted.

Oligo Sequences:

Human *SLC25A39* sg4 F: GATAGGCAGTGAAGTAGATGG;

sg4 R: CCATCTACTTCACTGCCTATC Human *GCLM* sg5 F: ACGGGGAACCTGCTGAACTG;

sg5 R: CAGTTCAGCAGGTTCCCCGT

sg7 F: TGGTGGCATCACACAGCAGG; sg7 R: CCTGCTGTGTGATGCCACCA

Primers targeting *GCLM* coding sequence: sg5 F: CTCCCTCTCGGGTCTCTCTC;

sg5 R: GACACTGGCCTCAGCATCT

sg7 F: TCACTGTGGATTGTGATTTTTCA; sg7 R: TGAGGGCATATAGGTTTTGAAC

### Generation of cDNA overexpression

As described in Wang et al.^28^, *S. thermophilus GshF* cDNA was humanized using the IDT codon-optimization tool and custom synthesized by IDT. To target GshF to mitochondria, the *Lactobacillus brevis COXIV* mitochondrial targeting sequence and human *ACO2* mitochondrial targeting sequence were added in tandem with a Flag-tag to the N-terminus of the humanized *GshF* cDNA.

Codon-optimized cDNA was cloned into BamH1, Not1-linearized pMXS-blasticidin or pMXS-blasticidin by Gibson reaction. Retroviral particles were produced by transfecting HEK 293F cells with cDNA-containing vector, viral packaging vector deltaVPR, and viral envelope vector gag-pol using XtremeGene 9 transfection reagent (Roche). Collection of viral particles and transduction was performed as described for lentiviral particles. Cells were selected with blasticidin or puromycin after removal of virus.

### cDNA Sequences

Below are *Homo sapiens* codon optimized sequences of *S. thermophilus* GshF with added mitochondrial targeting sequences (MTS) and FLAG-tags. To ensure mitochondrial localization of the mito-GshF construct, a tandem MTS consisting of the *L. brevis* COXIV MTS followed by the *H. sapiens* ACO2 MTS was added to the N-terminus of the GshF sequence, along with an N-terminal FLAG-tag. For clarity, corresponding regions of each sequence are highlighted as follows: FLAG tag, *L. brevis COXIV* MTS*, H. sapiens ACOII* MTS, (GGS)_3_ linker.

### Codon optimized *GshF*

ATGACCCTTAATCAGCTTCTCCAGAAGTTGGAGGCGACTTCCCCCATTCTCCAGGCGAACTTCGGGATAGAAAGGGAGTCATTGAGGGTTGACCGCCAGGGTCAGCTGGTCCACACACCGCACCCCTCATGTCTGGGAGCCCGCAGTTTTCATCCTTACATACAAACCGACTTTTGTGAATTCCAAATGGAACTGATTACACCAGTAGCCAAAAGTACGACGGAGGCCCGACGCTTTCTTGGCGCGATAACTGATGTAGCAGGACGAAGCATTGCAACTGACGAGGTGCTGTGGCCATTGAGTATGCCACCACGACTTAAAGCCGAGGAAATTCAAGTAGCGCAACTCGAGAACGACTTCGAAAGACATTATCGGAACTACTTGGCAGAGAAGTACGGCACCAAATTGCAGGCGATTAGTGGAATTCATTACAATATGGAACTTGGGAAGGACTTGGTTGAGGCGCTTTTTCAAGAGTCAGATCAGACTGACATGATCGCATTTAAAAACGCTCTGTATCTCAAGCTCGCCCAGAACTATTTGAGGTATCGGTGGGTCATTACTTACCTGTTTGGAGCAAGTCCCATTGCAGAACAAGGATTCTTTGACCAAGAAGTGCCGGAGCCTATGCGCTCTTTCCGCAACTCCGACCACGGCTACGTTAACAAGGAAGAGATACAGGTAAGCTTTGTATCCCTTGAAGACTATGTCTCCGCGATCGAGACCTACATCGAGCAGGGTGACCTTATAGCCGAGAAAGAGTTTTACTCAGCCGTGCGCTTTAGAGGACAAAAAGTCAATCGCTCCTTCCTTGATAAGGGTATAACTTATCTGGAGTTCAGAAACTTTGACTTGAACCCATTTGAGAGAATAGGCATCAGTCAAACCACTATGGATACCGTTCACTTGCTCATACTGGCCTTTCTCTGGTTGGATAGTCCGGAAAACGTGGACCAGGCCCTTGCCCAGGGACACGCGCTTAACGAGAAAATAGCCCTCTCCCATCCATTGGAGCCCCTCCCTTCAGAGGCCAAGACACAGGACATCGTGACCGCACTCGACCAGCTCGTACAGCACTTCGGATTGGGAGATTACCACCAGGATCTCGTGAAACAAGTGAAAGCGGCGTTTGCTGATCCGAATCAAACCCTGTCAGCTCAACTTCTTCCTTATATTAAGGACAAGTCACTCGCAGAATTCGCTCTCAATAAGGCACTCGCATATCATGACTATGACTGGACCGCTCACTACGCCCTTAAAGGTTACGAAGAAATGGAGCTCAGTACGCAGATGCTGCTCTTTGATGCTATCCAGAAAGGAATACACTTCGAGATACTCGATGAGCAAGATCAGTTCTTGAAGCTGTGGCATCAAGATCACGTAGAATATGTTAAAAACGGTAATATGACCAGCAAGGATAACTATGTAGTACCTCTCGCAATGGCCAACAAGACTGTTACTAAGAAAATTCTTGCTGACGCTGGGTTCCCTGTTCCGTCCGGGGACGAATTTACTAGCTTGGAGGAGGGACTGGCCTACTACCCGCTTATTAAAGATAAGCAAATT GTAGTAAAGCCAAAGAGCACGAATTTCGGCTTGGGTATCAGCATCTTCCAAGAACCCGCCAGTCTCGACAATTATCAAAAAGCATTGGAAATAGCATTTGCGGAGGACACTAGTGTGCTCGTCGAAGAATTCATTCCAGGCACGGAATACCGATTCTTCATTTTGGACGGACGCTGTGAGGCAGTCCTTTTGAGGGTAGCTGCCAATGTAATAGGGGACGGGAAACACACAATCAGAGAGTTGGTAGCGCAGAAAAACGCAAATCCCCTGCGCGGTAGGGATCATAGATCACCCTTGGAAATCATAGAGCTTGGGGACATAGAGCAACTCATGCTGGCACAGCAGGGTTACACTCCAGATGACATCCTGCCAGAGGGTAAGAAAGTGAATTTGAGGCGGAACAGCAATATTAGTACTGGGGGAGACTCCATAGACGTCACAGAAACAATGGATAGCTCTTATCAAGAACTTGCAGCAGCGATGGCTACCAGTATGGGGGCATGGGCCTGTGGAGTTGATCTGATTATACCCGACGAAACGCAGATTGCCACAAAGGAAAATCCACATTGCACGTGTATTGAACTTAACTTCAACCCCTCCATGTACATGCATACATACTGCGCTGAGGGGCCGGGGCAGGCAATTACAACCAAAATACTCGACAAACTCTTCCCGGAGATCGTGGCCGGACAAACTGGAGGAAGCGGAGGAAGCGGAGGAAGCGATTACAAGGATGACGATGACAAGTAA

### Mito-GshF sequence

ATGCTCGCTACAAGGGTCTTTAGCCTCGTCGGAAAGAGAGCTATCAGCACCTCCGTCTGCGTGAGAGCTCATatggcgccctacagcctactggtgactcggctgcagaaagctctgggtgtgcggcagtaccatgtggcctcagtcctgtgcGATTACAAGGATGACGATGACAAGGGAGGAAGCGGAGGAAGCGGAGGAAGCACCCTTAATCAGCTTCTCCAGAAGTTGGAGGCGACTTCCCCCATTCTCCAGGCGAACTTCGGGATAGAAAGGGAGTCATTGAGGGTTGACCGCCAGGGTCAGCTGGTCCACACACCGCACCCCTCATGTCTGGGAGCCCGCAGTTTTCATCCTTACATACAAACCGACTTTTGTGAATTCCAAATGGAACTGATTACACCAGTAGCCAAAAGTACGACGGAGGCCCGACGCTTTCTTGGCGCGATAACTGATGTAGCAGGACGAAGCATTGCAACTGACGAGGTGCTGTGGCCATTGAGTATGCCACCACGACTTAAAGCCGAGGAAATTCAAGTAGCGCAACTCGAGAACGACTTCGAAAGACATTATCGGAACTACTTGGCAGAGAAGTACGGCACCAAATTGCAGGCGATTAGTGGAATTCATTACAATATGGAACTTGGGAAGGACTTGGTTGAGGCGCTTTTTCAAGAGTCAGATCAGACTGACATGATCGCATTTAAAAACGCTCTGTATCTCAAGCTCGCCCAGAACTATTTGAGGTATCGGTGGGTCATTACTTACCTGTTTGGAGCAAGTCCCATTGCAGAACAAGGATTCTTTGACCAAGAAGTGCCGGAGCCTATGCGCTCTTTCCGCAACTCCGACCACGGCTACGTTAACAAGGAAGAGATACAGGTAAGCTTTGTATCCCTTGAAGACTATGTCTCCGCGATCGAGACCTACATCGAGCAGGGTGACCTTATAGCCGAGAAAGAGTTTTACTCAGCCGTGCGCTTTAGAGGACAAAAAGTCAATCGCTCCTTCCTTGATAAGGGTATAACTTATCTGGAGTTCAGAAACTTTGACTTGAACCCATTTGAGAGAATAGGCATCAGTCAAACCACTATGGATACCGTTCACTTGCTCATACTGGCCTTTCTCTGGTTGGATAGTCCGGAAAACGTGGACCAGGCCCTTGCCCAGGGACACGCGCTTAACGAGAAAATAGCCCTCTCCCATCCATTGGAGCCCCTCCCTTCAGAGGCCAAGACACAGGACATCGTGACCGCACTCGACCAGCTCGTACAGCACTTCGGATTGGGAGATTACCACCAGGATCTCGTGAAACAAGTGAAAGCGGCGTTTGCTGATCCGAATCAAACCCTGTCAGCTCAACTTCTTCCTTATATTAAGGACAAGTCACTCGCAGAATTCGCTCTCAATAAGGCACTCGCATATCATGACTATGACTGGACCGCTCACTACGCCCTTAAAGGTTACGAAGAAATGGAGCTCAGTACGCAGATGCTGCTCTTTGATGCTATCCAGAAAGGAATACACTTCGAGATACTCGATGAGCAAGATCAGTTCTTGAAGCTGTGGCATCAAGATCACGTAGAATATGTTAAAAACGGTAATATGACCAGCAAGGATAACTATGTAGTACCTCTCGCAATGGCCAACAAGACTGTTACTAAGAAAATTCTTGCTGACGCTGGGTTCCCTGTTCCGTCCGGGGACGAATTTACTAGCTTGGAGGAGGGACTGGCCTACTACCCGCTTATTAAAGATAAGCAAATTGTAGTAAAGCCAAAGAGCACGAATTTCGGCTTGGGTATCAGCATCTTCCAAGAACCCGCCAGTCTCGACAATTATCAAAAAGCATTGGAAATAGCATTTGCGGAGGACACTAGTGTGCTCGTCGAAGAATTCATTCCAGGCACGGAATACCGATTCTTCATTTTGGACGGACGCTGTGAGGCAGTCCTTTTGAGGGTAGCTGCCAATGTAATAGGGGACGGGAAACACACAATCAGAGAGTTGGTAGCGCAGAAAAACGCAAATCCCCTGCGCGGTAGGGATCATAGATCACCCTTGGAAATCATAGAGCTTGGGGACATAGAGCAACTCATGCTGGCACAGCAGGGTTACACTCCAGATGACATCCTGCCAGAGGGTAAGAAAGTGAATTTGAGGCGGAACAGCAATATTAGTACTGGGGGAGACTCCATAGACGTCACAGAAACAATGGATAGCTCTTATCAAGAACTTGCAGCAGCGATGGCTACCAGTATGGGGGCATGGGCCTGTGGAGTTGATCTGATTATACCCGACGAAACGCAGATTGCCACAAAGGAAAATCCACATTGCACGTGTATTGAACTTAACTTCAACCCCTCCATGTACATGCATACATACTGCGCTGAGGGGCCGGGGCAGGCAATTACAACCAAAATACTCGACAAACTCTTCCCGGAGATCGTGGCCGGACAAACTTAA

### Codon-optimized *SLC25A39* sequence

ATGGCAGACCAGGACCCCGCGGGCATCTCACCTCTCCAGCAGATGGTCGCATCTGGAACAGGGGCAGTCGTCACAAGTTTGTTCATGACCCCACTTGATGTAGTGAAAGTCCGGCTTCAATCACAACGCCCTAGCATGGCCAGCGAGCTGATGCCGAGCTCCAGGCTCTGGTCACTTTCTTATACGAAGCTTCCCTCTTCTCTCCAGTCTACGGGTAAATGTTTGCTTTATTGTAACGGCGTACTCGAACCTCTGTATTTGTGTCCAAATGGAGCACGCTGCGCCACGTGGTTTCAGGACCCAACTCGATTTACCGGCACAATGGACGCATTTGTCAAGATAGTAAGACACGAGGGTACAAGAACGCTTTGGAGCGGCCTCCCTGCTACGTTGGTGATGACGGTTCCCGCAACGGCCATATACTTTACAGCCTACGACCAGCTGAAGGCCTTTCTGTGTGGTAGGGCACTTACCTCAGACCTTTACGCTCCAATGGTCGCAGGGGCCCTTGCAAGACTTGGTACGGTCACTGTAATAAGTCCGCTCGAACTCATGAGGACAAAACTCCAAGCTCAGCACGTGAGCTACCGGGAACTGGGGGCTTGTGTACGCACAGCGGTCGCGCAAGGCGGCTGGAGGAGTCTGTGGCTGGGTTGGGGGCCCACGGCCCTCCGGGACGTACCGTTTTCTGCGCTTTATTGGTTTAACTACGAGCTTGTGAAATCTTGGCTCAATGGATTCCGGCCGAAAGACCAGACCTCCGTTGGAATGTCTTTCGTCGCCGGGGGCATTTCCGGCACGGTGGCCGCCGTGCTGACCTTGCCATTCGACGTTGTTAAGACCCAGCGACAGGTCGCTTTGGGGGCAATGGAGGCCGTGCGGGTGAACCCACTCCACGTTGACAGTACATGGTTGCTGCTCCGCCGCATCCGGGCCGAAAGCGGAACTAAAGGTCTGTTTGCTGGATTTCTTCCGCGAATCATTAAGGCTGCGCCATCTTGTGCAATCATGATCTCTACATACGAGTTTGGAAAATCCTTCTTTCAGAGGCTTAATCAGGACAGACTGCTCGGAGGGTAA

### Codon-optimized *Gclm* sequence

ATGGGCACCGACAGCCGCGCGGCCAAGGCGCTCCTGGCGCGGGCCCGCACCCTGCACCTGCAGACGGGGAACCTGCTGAACTGGGGCCGCCTGCGGAAGAAGTGCCCGTCCACGCACAGCGAGGAGCTTCATGATTGTATCCAAAAAACCTTGAATGAATGGAGTTCCCAAATCAACCCAGATTTGGTCAGGGAGTTTCCAGATGTCTTGGAATGCACTGTATCTCATGCAGTAGAAAAGATAAATCCTGATGAAAGAGAAGAAATGAAAGTTTCTGCAAAACTGTTCATTGTAGAATCAAACTCTTCATCATCAACTAGAAGTGCAGTTGACATGGCCTGTTCAGTCCTTGGAGTTGCACAGCTGGATTCTGTGATCATTGCTTCACCTCCTATTGAAGATGGAGTTAATCTTTCCTTGGAGCATTTACAGCCTTACTGGGAGGAATTAGAAAACTTAGTTCAGAGCAAAAAGATTGTTGCCATAGGTACCTCTGATCTAGACAAAACACAGTTGGAACAGCTGTATCAGTGGGCACAGGTAAAACCAAATAGTAACCAAGTTAATCTTGCCTCCTGCTGTGTGATGCCACCAGATTTGACTGCATTTGCTAAACAATTTGACATACAGCTGTTGACTCACAATGATCCAAAAGAACTGCTTTCTGAAGCAAGTTTCCAAGAAGCTCTTCAGGAAAGCATTCCTGACATTCAAGCGCACGAGTGGGTGCCGCTGTGGCTACTGCGGTATTCGGTCATTGTGAAAAGTAGAGGAATTATCAAATCAAAAGGCTACATTTTACAAGCTAAAAGAAGGGGTTCTTAA

### Cell Fractionation

Cell fractionation was performed according to Cell Fractionation Kit (9038, Cell Signaling Technology) instructions, with the following modifications: 2.5e6 cells were pelleted at 350xg for 5 min, washed twice with ice-cold PBS, and resuspended in 500μL cold PBS. 100μL of cell suspension was collected for the whole cell lysate (WCL). Remaining cells were pelleted at 500xg at 4°C, and resuspended in 250μL Cytoplasm Isolation Buffer (CIB) containing protease inhibitors (Sigma-Aldrich, 11836170001). After saving the supernatant, the pellet was washed 3x with 100μL CIB before resuspension in 250μL Membrane Isolation Buffer (MIB) containing protease inhibitor. After removing the membrane and organelle fraction, the pellet was washed 3X with 100μL MIB before resuspending the cytoskeletal and nuclear fraction in 125 μL Cytoskeletal and Nuclear Isolation Buffer. Samples were boiled at 95°C and centrifuged for 3min at 15,000 x g before resolution on 10-20% SDS-PAGE gel.

### Immunoblotting

Cell pellets were washed twice with ice-cold PBS before lysis in ice-cold RIPA buffer (10mM Tris-HCl, pH 7.5, 150mM NaCl, 1mM EDTA, 1% Triton X-100, 0.1% SDS) supplemented with protease inhibitors (Sigma-Aldrich, 11836170001) and phosphatase inhibitors (Roche, 04906837001). Lysis was allowed to proceed on ice for 30 minutes before centrifugation at 20,000xg for 10 min at 4°C. Supernatant was collected and protein concentration was determined by BCA (Thermo Scientific, 23227). Except for cell fractionation, samples were diluted to equal concentrations and heated at 70°C for 10 min before resolution on 12% or 10-20% SDS-PAGE gels (Invitrogen). Resolved proteins were transferred to nitrocellulose membrane in CAPS transfer buffer. Membranes were blocked in 5% BSA in 1X TBST before incubation in primary antibodies at 4°C overnight with shaking. Membranes were incubated with HRP-linked anti-mouse IgG (Cell Signaling, 7076, 1:5000) or anti-rabbit IgG (Cell Signaling, 7074, 1:5000) for 1hr at room temperature while shaking. Membranes were incubated in PerkinElmer Enhanced Chemiluminescence Substrate (PerkinElmer, NEL105001EA) for 1 minute. HyBlot CL Autoradiography Film (Thomas Scientific, 114J52) was exposed to membranes in the dark and developed on an SRX-101A Film Processor (Konica Minolta).

For tissue samples, a 4mm biopsy punch from snap-frozen tissue was homogenized in 500μL RIPA buffer with protease and phosphatase inhibitors in a 2mL dounce homogenizer. Samples were rotated at 4°C for 10 minutes before centrifugation for 2 minutes at 1000xg at 4°C to remove excess tissue. 10μL of the supernatant was reserved to determine protein concentration, and Laemmli dye was added to 150μL of supernatant. Samples were diluted to equal concentrations and sonicated before Western blots were performed as described above.

### Polar Metabolite Profiling

For whole cell samples, cells were washed twice with 1mL ice-cold 0.9% NaCl. Polar metabolites were extracted in 500μL ice-cold 80% methanol containing amino acid standards (Cambridge Isotope Labs, MSK-A2-1.2). Samples were vortexed at 4°C for 10 minutes and centrifuged for 10 minutes at 20000xg at 4°C before supernatant was collected and dried via nitrogen evaporation. Total protein in the pellet was determined by BCA to normalize metabolite levels after quantification by mass spectrometry. Dried samples were stored at -80°C until resuspension in 60μL of 50% acetonitrile for metabolomics analysis.

Mitochondria were purified from HEK 293T cells expressing 3xHA-OMP25-mCherry or a negative control expressing 3xMyc-OMP25-mCherry according to the protocol described by Chen et al.^63^ In brief, triplicate samples of 25-30 million cells were collected per condition. Cells were washed twice with ice-cold 0.9% NaCl and scraped into 1mL of ice-cold KPBS. Cells were spun at 1000xg for 1.5 min at 4°C and resuspended in 1mL of cold KPBS. 10μL of resuspended cells were transferred for input protein and whole cell metabolites were extracted from an additional 10μL transferred into 40μL of 80% methanol with amino acid standards. The remaining sample was transferred to a 2mL dounce homogenizer and homogenized with two sets of 30 strokes. The homogenate was centrifuged and supernatant was incubated with 200μL of 50% anti-HA magnetic beads (Thermo Scientific Pierce 88837) in KPBS for 5 minutes while rotating at 4°C. Beads were washed three times in 1mL cold KPBS. 10% of bead volume was taken for protein and metabolites were extracted from the remaining bead volume with 50μL of 80% methanol containing amino acid standards. Samples were rotated at 4°C for 10 minutes before being spun at 20000xg for 10 minutes at 4°C. Input and mitochondrial samples were subjected to LC-MS profiling without drying. Data were normalized to citrate synthase or GAPDH protein level as quantified from Western blot or NAD^+^ abundance.

Tissue samples were immediately flash frozen in liquid nitrogen after rapid isolation from mice euthanized by cervical dislocation. Samples were stored at -80°C until metabolite extraction. To extract metabolites, 2x 4mm biopsy punches were taken from snap-frozen tissue and homogenized in 1mL ice cold 0.9% NaCl in a 2mL dounce homogenizer with 2 rounds of 35 strokes. Excess tissue was removed by centrifugation at 1000xg for 2 min at 4° C. Metabolites were extracted from 10μL of supernatant with 40μL of 1:1 acetonitrile/methanol containing amino acid standards. Samples were vortexed for 10 minutes at 4°C before being centrifuged at 20000xg for 10 min at 4°C. The supernatant was dried and stored at -80°C until resuspension in 2:2:1 acetonitrile/methanol/water for mass spectrometry analysis.

For plasma samples, ∼100uL whole blood was collected into EDTA-coated tubes containing 20μL of N-ethylmaleimide (NEM), amino acid standards, and 5μL of 1mM [^12^C_2_, ^15^N]-GSH standard. Samples were vortexed briefly on ice before centrifugation twice at 400xg for 5min to obtain plasma. Dithiothreitol (DTT, Sigma) was then added to equal the final concentration of NEM in plasma. Proteins were precipitated with 80% methanol for 30 minutes at -20°C and removed by centrifugation at 20000xg for 10 minutes. The supernatant was dried under nitrogen and resuspended in 100μL of cold water, and the pellets were reserved to determine total protein by BCA. The samples were vortexed for 10 seconds before addition of 300μL cold dichloromethane. Samples were vortexed vigorously for 10 minutes at 4°C prior to centrifugation at 20000xg for 10 minutes. The polar phase was dried and stored at -80°C for no more than 48 hours before resuspension in 50μL of 75:25 acetonitrile/methanol for metabolomics analysis.

Metabolomics analysis was performed on a QExactive benchtop orbitrap mass spectrometer equipped with an IonMax source and a HESI II probe coupled to a Dionex UltiMate 3000 UPLC system (Thermo Fisher Scientific). External mass calibration was performed using the standard calibration mixture every 7 days. Two microliters of sample were injected into a ZIC-pHILIC 150 x 2.1 mm (5μm particle size) column (EMD Millipore). Chromatographic separation was achieved using the following conditions: buffer A was 20mM ammonium carbonate, 0.1% ammonium hydroxide; buffer B was acetonitrile. The column oven and autosampler tray were held at 25°C and 4°C respectively. The chromatographic gradient was run at a flow rate of 0.15 mL/min as follows: 0-20 min linear gradient from 80% to 20% B; 20-20.5 min linear gradient from 20% 80% B; 20.5-28 min hold at 80% B. The mass spectrometer was operated in full-scan, polarity switching mode with the spray voltage set to 3.0kV, the heated capillary held at 275°C, and the HESI probe held at 350°C. The sheath gas flow was set to 40 units, the auxiliary gas flow was set to 15 units, and the sweep gas flow was set to 1 unit. The MS data acquisition was performed in a range of 70-1000 *m/z*, with the resolution set at 70,000, the AGC target at 10e6, and the maximum injection time at 20ms. Relative quantification of polar metabolites was performed with XCalibur QuanBrowser 2.2 using a 5ppm mass tolerance and referencing an in-house library of chemical standards. Metabolite levels were normalized to the total protein amount for each sample, determined by BCA on the pellet of each sample.

### Metabolite Tracing

For cystine tracing experiment, HEK 293T *Gclm*-knockout cells expressing indicated cDNAs were seeded in triplicate in 6 well plates. Cells were allowed to attach for 24 hours prior to addition of BSO. After 24 hours of BSO treatment, media was replaced with RPMI + 10% dialyzed FBS (Gibco) lacking cystine and containing [^13^C_6_, ^15^N_2_]-L-Cystine (207.67μM). After 24 hours of incubation with [^13^C_6_, ^15^N_2_]-L-Cystine in the presence or absence of BSO, metabolites were extracted and quantified as described above. Fractional labeling was corrected for natural abundance with IsoCorrectoR^64^ using RStudio version (2022.12.0+353).

### Proliferation Assays

Cells were seeded in triplicate in 96-well plates as follows: Jurkat cells, 2000 cells/well; PANC-1 and HepG2, 500 cells/well; all other cell lines 1000 cells/well. Cells were allowed to attach to plates for 6 hours prior to addition of compounds described in each experiment. An additional plate was seeded without treatment and taken for an initial time point upon addition of compounds to the experimental plates. Cells were allowed to proliferate for 5 days. At either the initial timepoint or the endpoint, 40μL of Cell Titer Glo reagent (Promega) was added to each well. Plates were incubated in the dark on a rocker for 10 minutes prior to quantification of luminescence per well on a SpectraMax M3 plate reader (Molecular Devices). The fold change in luminescence relative to day 0 was determined and reported on a log_2_ scale.

### CRISPR Screens

The metabolism-focused human sgRNA library used was first described by Birsoy et al.,^65^ and screens were performed as described previously.^65–67^ Olignonucleotides for sgRNAs were synthesized by CustomArray and amplified by PCR. In brief, a lentiviral library was generated from the plasmid pool and transfected into HEK 293F cells to produce viral particles as described above. *SLC25A39*-knockout Jurkat cells expressing either *GshF* or *mito-GshF* were infected at a multiplicity of infection of 0.7 and selected with puromycin for 3 days. An initial sample of 30 million cells was taken for each cell line, and cells were seeded into 500mL spinning flasks in RPMI alone or RPMI with 200μM BSO (Sigma). Cells were split every 3 days and grown under indicated conditions for approximately 14 population doublings. Final samples of 30 million cells were collected, and DNA was extracted according to DNeasy Blood and Tissue Kit instructions (Qiagen 69506). sgRNA inserts were amplified and barcoded by PCR with primers unique to each condition. PCR amplicons were gel-purified and sequenced on a NextSeq500 (Illumina). Screens were analyzed using Python (v.2.7.13), R (v.3.3.2), and Unix (v.4.10.0-37-generic x86_64). The gene score for each gene was defined as the median log2 fold change in the abundance of each sgRNA targeting that gene. A complete list of differential gene scores for each screen is provided in Supporting Table S1.

### Generation of transgenic mouse strains

All animal studies and procedures were conducted according to a protocol approved by the Institutional Animal Care and use Committee (IACUC) at the Rockefeller University. All mice were maintained on a standard light:dark cycle with food and water provided ad libitum.

*GshF* cDNA was cloned into FseI-linearized Ai9 vector, which was previously reported to result in broad expression patterns when inserted at the Rosa26 locus.^68^ *mito-GshF* cDNA was cloned into a CTV vector. Each of these targeting constructs contains 5’ and 3’ *Rosa26* homology arms and a LoxP-STOP-LoxP cassette upstream of the inserted cDNA. Each targeting construct was electroporated into Cy2.4 [B6(Cg)-Tyr <C2J> genetic background] embryonic stem (ES) cells. cDNA-containing ES clones were selected with neomycin, and presence of the cDNA was validated by PCR and with anti-Flag immunoblots from Cre-transfected clones. Two single clones for each construct were injected into blastocysts to produce germline-transformed heterozygous GshF and mito-GshF mice. Chimeras were screened by coat color for back-crossing to wild-type C57BL6/J (The Jackson Laboratory, 000664) and genotyped for subsequent matings. All mice were backcrossed for at least 4 generations prior to conducting any experiments shown here. GshF and mito-GshF mice were generated in the Birsoy laboratory with the assistance of the CRISPR and Genome Editing Center at the Rockefeller University.

CMV-Cre (006054) and ER^T^^2^-Cre (008463) mice were purchased from The Jackson Laboratory.

### Mouse strain accessibility

GshF^fl/fl^ and mito-GshF^fl/fl^ have been deposited to the Jackson Laboratory under stock numbers 038275 and 038277, respectively.

### Mouse Studies

All mice were intraperitonially injected with 100μL tamoxifen (Cayman Chemical, 13258) on alternating abdominal sides for 5 consecutive days. Tamoxifen was prepared at 20mg/mL in sterile corn oil (Sigma C8267) and shaken at 32°C until in solution. All experiments were performed at least 7 days after the final tamoxifen injection. All mice used in these studies were 8-14 weeks old, and mice of both sexes were used except when specified in figure legends. Both ER^T2^-Cre -null, GshF-positive and ER^T2^-Cre-positive, GshF-null mice were used as GshF-null controls.

### Genotyping

Mito-GshF^fl/fl^ and CMV-mito-GshF mice were genotyped by Transetyx. GshF^fl/fl^, CMV-GshF, and ERT2-GshF mice were genotyped by Transnetyx and/or using the following primers:

*GshF* F: CCGCGGGCCCTAAGAAGTTCCT;

R: CCATTTGGAATTCACAAAAGTCGGTTTGT

*Cre* F: GCGGTCTGGCAGTAAAAACTATC; R: GTGAAACAGCATTGCTGTCACTT

*Rosa26* F: AGCCTTTAAGCCTGCCCAGAAGACTCC;

R: TGCTCTCCCAAAGTCGCTCTGAGTTGTTAT

*GshF* primers result in a ∼900 bp band when floxed and a ∼370bp band when induced by Cre. *Cre* primers only produce bands when *Cre* is present; *Rosa26* primers only produce bands when the locus is wild-type.

### Embryo Imaging

Timed pregnancies were determined by confirmation of a visible vaginal plug the morning after mating pairs were combined. After confirmation of vaginal plug, males were removed from cages to prevent additional copulation. The day that vaginal plugs are identified is considered gestational day E0.5. Females were euthanized by CO_2_ exposure at indicated timepoints and embryos were rapidly collected. For embryos younger than E15.5, egg sacs were collected for genotyping; for E15.5 and E17.5 tail clippings were collected for genotyping. Brightfield images were taken with a Zeiss Axiocam 506 mono camera on an AxioZoom V.16 microscope. Embryos were fixed in formalin for 48 hours. Sectioning and H&E staining was performed by Histoserv. Sagittal sections were taken at the midline, at the kidney, and at the eye level. Histopathology evaluation was performed by Ileana Miranda at the Laboratory of Comparative Pathology at Memorial Sloan Kettering.

### Complete Blood Counts

Whole blood was collected in EDTA-coated tubes. Complete blood counts were performed within 30 minutes of blood sampling on a Heska Element HT5 hematology analyzer.

### Immune Profiling

Mice were injected retro-orbitally with 2μg/100uL CD45-FITC antibody (BioLegend) in PBS. Three minutes post injection, mice were sacrificed by cervical dislocation. Mesenteric lymph nodes, spleen, thymus, lung, and colon were rapidly removed for processing.

For mesenteric lymph nodes, spleen, and thymus, tissues were ground in a 6-well plate through a 70μm cell strainer (Corning 352350) in T Cell Media (TCM) consisting of RPMI 1640 (Gibco 21870) supplemented with 10% FBS (Gibco 10437), 1% Penicillin-Streptomycin (Gibco 15140), 1% L-glutamine (Gibco 25030), 1% sodium pyruvate (Gibco 113690), 1% Non-Essential amino acids (Gibco 11140), 1% HEPES (Gibco 15630), and 50μM β-mercaptoethanol (Sigma M6250). Cells were transferred to 15mL conical tubes and centrifuged. Spleen cells were resuspended in red blood cell lysis buffer (Sigma 11814389001), incubated at room temperature for 3 minutes, and washed with TCM before staining. Lungs were washed in cold PBS and transferred to gentleMACS™ C Tubes (Miltenyi Biotec 130-096-334) containing 3mL of TCM. For digestion, 75μL of Collagenase A (Sigma 10103578001, stock at 20mg/mL) and 10μL DNase (Qiagen, stock at 10mg/mL) were added to each tube, and processed in gentleMACS Dissociator (Miltenyi Biotec 130-093-235), program Lung_LIDK_37. Lungs were further ground and filtered through a 70μm cell strainer. Cells were washed with TCM, centrifuged, and resuspended in 7mL of 40% Percoll (GE Healthcare 17-0891-02). 1.5mL of 80% Percoll was underlaid using a Pasteur pipette (Fisher 13-678-20D). Cells were centrifuged at 2300rpm for 25 minutes at 20°C, with brakes off. Tubes were carefully removed not to disturb the gradient, and ∼ 5mL were aspirated without disturbing the cells in the interface between 40% and 80% Percoll. Cells were washed with TCM and inverted multiple times to mix before centrifugation and collection for staining.

Colons were processed as previously described for the small intestine.^69^ Briefly, colons were filet-opened, washed with cold PBS, chopped into 50 mL conical tubes filled with 10mL of PBS and supplemented with 1μM DTT (Sigma). Tubes were shaken at room temperature for 10 minutes, then intensely shaken by hand for 2 minutes before tissue was filtered through a metal strainer into a new 50mL conical tube containing with 25mL of cRPMI (RPMI Media 1640 supplemented with 1% HEPES and 2% FCS). Flowthrough was considered Fraction 1 of intraepithelial lymphocytes (IEL). Tissues remaining in the strainer were transferred to 10mL of PBS containing 30mM EDTA (ThermoFisher Scientific) and 10mM HEPES, and tubes were shaken in a 37°C shaker at 180rpm for 10 minutes. Tubes were then shaken vigorously by hand for 2 minutes, and supernatant filtered into IEL Fraction 1. The combined IEL fractions were filtered in a 40% to 80% Percoll gradient as previously described. Remaining tissues were transferred to 10 mL of PBS containing 30mM EDTA and 10mM HEPES and shaken at 180 rpm for 10 minutes at 37°C followed by 2 minutes of vigorous hand shaking. The supernatant was discarded and tissues were washed several times with cold PBS before being finely minced with scissors in 6-well plates. The tissue slurry was resuspended in 6mL of collagenase mix (Collagenase 8 and DNase in TCM) and while shaken for 45 minutes at 37°C, 80rpm before filtration through 70μm mesh. Samples were then washed and filtered through a Percoll gradient as previously described. All centrifugations were done at 400xg for 5 minutes at 4°C, except for the Percoll gradient.

For flow cytometry analysis, cells were stained in a surface antibody mix in PBS for 25 minutes at 4°C, washed in PBS, then fixed/ permeabilized for 1hr at 4°C using eBioscience™ Foxp3/ Transcription Factor Staining Buffer Set (Thermo Fisher Scientific). Cells were then washed with 1X permeabilization buffer (Thermo Fisher Scientific) and stained in intracellular antibody mix for 45-90 minutes at 4°C and washed with PBS before flow cytometry analysis.

### RNAseq

Tissues were rapidly isolated and snap-frozen in liquid nitrogen from mice euthanized by cervical dislocation. RNA was extracted from whole kidney or single lung using TRIzol reagent (Thermo Fisher Scientific) according to manufacturer’s manual. After DNase I treatment (New England Biolabs), RNA concentrations were determined using a Qubit 2.0 Fluorometer (Life Technologies) and RNA integrity was confirmed by Agilent TapeStation 4200 (Agilent Technologies). RNA sequencing libraries were prepared with unique barcodes using the NEBNext Ultra RNA library kit for Illumina (New England Biolabs) according to manufacturer’s instructions. The sequencing libraries were pooled at equimolar ratios and sequenced on an Illumina NextSeq 500 using 2×150bp paired-end reads. Read quality was confirmed with Rfastp.^70^ The reads alignment and gene models were based on Mus_musculus.GRCm39.107.gtf.gz. Transcript expression was computed using Salmon^71^, followed by processing with DESeq2.^72^ Significant differentially expressed genes between conditions were identified by DESeq2 with a Benjamini-Hochberg adjusted p-value < 0.05 and absolute fold change > 0.5. All RNAseq results are available in Supporting Tables S2 and S3. Gene Ontology (GO) terms were retrieved from Mouse MSigDB Collections Gene Ontology Analysis.^73–75^ Terms consisting of fewer than 10 or more than 500 were excluded. The GO analysis was performed by using the hypergeometric test followed by Benjamini-Hochberg multiple test correction.

### Statistics and Reproducibility

GraphPad PRISM 9 and 10 and Microsoft Excel Software (version 16.77.1) were used for statistical analysis. Skyline-daily 64-bit version 21.2.1455 was used for metabolomics analysis, and FIJI (ImageJ2, NIH, version 2.14.0/1.54f) was used for image analysis including measuring embryo length. Error bars and statistical tests are reported in the figure captions. For all figures, only *P* values less than or equal to 0.05 are reported unless otherwise noted. All experiments except RNAseq and CRISPR screens were performed at least twice with similar results. Both technical and biological replicates were reliably reproduced. Comparison of two mean values was evaluate by two-tailed unpaired *t-*test. Comparison of multiple mean values was evaluated by one-way ANOVA followed by post hoc Bonferroni test. Comparison of multiple mean values under different conditions was evaluated by two-way ANOVA.

## Data Availability

All data supporting the findings in this study are available upon request from the corresponding author.

## Acknowledgements

We thank all members of the Birsoy lab for helpful suggestions. Additionally, we’d like to thank Ileana Miranda from the Memorial Sloan Kettering Laboratory of Comparative Pathology for the pathology analysis and Rada Norinsky from the Rockefeller University Transgenic Services department for performing the blastocyst injections as well as *in vitro* fertilization to expand the mouse colony. K.B. was supported by NCI (R01CA273233) and Mark Foundation Emerging Leader Award; and is a Searle and Pew-Stewart Scholar. RCT was supported by Agnes Gund through the Anna-Maria and Stephen Kellen Women’s Entrepreneurship Fund and the Women & Science Women’s Entrepreneurship Fund.

